# Cryo-EM captures the coordination of long-range allostery and asymmetric electron transfer by a bi-copper cluster in the nitrogenase-like DPOR complex

**DOI:** 10.1101/2024.04.26.590571

**Authors:** Rajnandani Kashyap, Jaigeeth Deveryshetty, Natalie Walsh, Monika Tokmina-Lukaszewska, Brian Bothner, Brian Bennett, Edwin Antony

## Abstract

Enzymes that catalyze long-range electron transfer reactions are often structurally evolved to possess two symmetrical halves. The functional advantages and mechanistic principles for such architecture remain a mystery. Using Cryo-EM we capture snapshots of the nitrogenase-like Dark-operative Protochlorophyllide Oxidoreductase (DPOR) enzyme during substrate recognition and turnover. The structures reveal that asymmetry is enforced upon substrate binding and leads to an allosteric inhibition of protein-protein interactions and electron transfer in one half. Residues that form a conduit for electron transfer are aligned in one half while misaligned in the other. An ATP-turnover coupled switch is triggered once electron transfer is accomplished in one half and relayed through a bi-copper cluster at the oligomeric interface, leading to activation of enzymatic events in the other. The findings provide a mechanistic blueprint for regulation of asymmetric long-range electron transfer.

**One-Sentence Summary:** A bi-copper cluster coordinates electron transfer for substrate reduction in the nitrogenase-like DPOR enzyme and the structures reveal how allostery and asymmetry are enacted over 100Å and utilized for sequential electron transfer.

---

Photosynthetic organisms employ two distinct approaches for reducing protochlorophyllide (Pchlide) to chlorophyllide (Chlide), a key intermediate in the chlorophyll biosynthetic pathway(*1*). In angiosperms, this reductive step is catalyzed by light dependent Pchlide oxidoreductase (LPOR)(*2*). In photosynthetic bacteria, cyanobacteria, green algae, and gymnosperms this reaction is catalyzed in the dark by the ATP-dependent dark-operative protochlorophyllide oxidoreductase (DPOR)(*3*). While these enzymes are not structurally related, they both catalyze the 2-electron reduction of the C17=C18 double bond in Pchlide using differing mechanisms (**Fig. 1A**)(*4*). DPOR shares structural homology with nitrogenase, the enzyme that catalyzes the reduction of dinitrogen to ammonia(*5, 6*), and thus is classified as a nitrogenase-like enzyme. Nitrogenase, DPOR, and other nitrogenase-like enzymes are composed of electron donor and electron acceptor component proteins(*7, 8*). In DPOR, the homodimeric BchL is the donor (L-protein) with an ATPase site per subunit and one [4Fe-4S]^L^ cluster coordinated at the dimer interface (**Fig. 1B**)(*9*). The acceptor, BchN-BchB, is an α_2_β_2_ heterotetramer (NB-protein) with two structurally identical halves (**Fig. 1B**)(*10*). Each half possesses a [4Fe-4S]^NB^ cluster and an active site for Pchlide binding and reduction. ATP binding to the L-protein promotes transient association of the component proteins resulting in one electron being transferred to the substrate (Pchlide)(*11*). Two such rounds of electron transfer (ET) are required to reduce Pchlide to Chlide.

**Figure 1.**
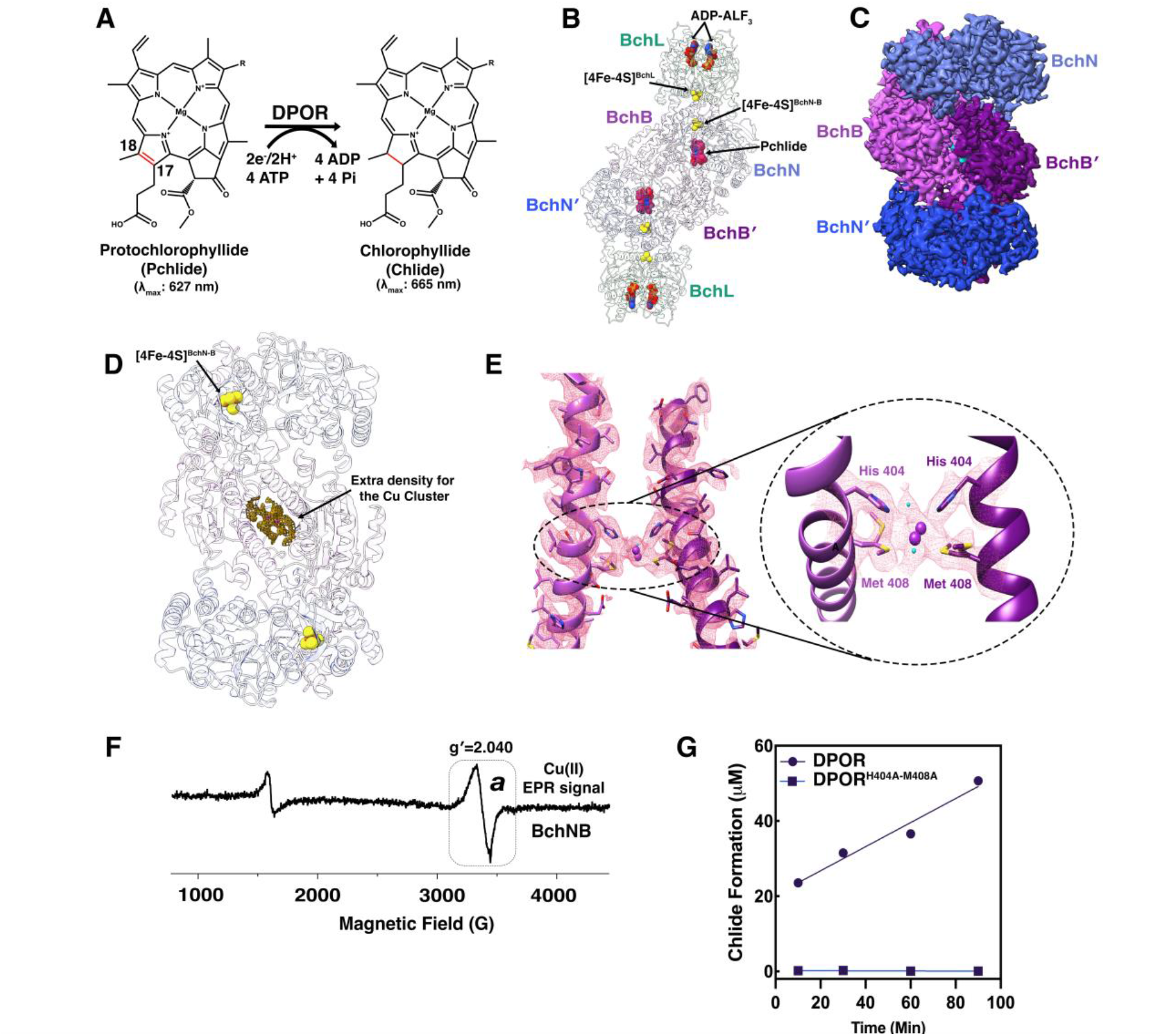
NB-protein possesses a bi-copper cluster. (**A**) Schematic of the chemical reaction catalyzed by DPOR. Two rounds of electron transfer are required to reduce the C17=C18 double bond (shown in red) of Pchlide to form Chlide. (**B**) Crystal structure of *P. marinus* DPOR (PDB: 2YNM) stabilized in the presence of ADP-AlF_3_ depicting symmetrical binding of two L-protein homodimers to a NB-protein heterotetramer. The [4Fe-4S]^L^ and [4Fe-4S]^NB^ clusters and the Pchlide molecules are highlighted. (**C**) A 2.7Å Cryo-EM map of *R. sphaeroides* NB-protein. The subunits are colored light blue (BchN), dark blue (BchN’), light purple (BchB), and dark purple (BchB’), respectively. (**D**) Representative structure of the NB-protein showing the position of [4Fe-4S]^NB^ cluster and the extra density for the bi Cu cluster at the interface. (**E**) Carved electron density for the two helices coordinating the Cu cluster and the His-404 and Met-408 residues that interact with the cluster at the NB-protein tetramer interface. Water molecules are colored cyan. (**F**) EPR spectra of the NB-protein shows a signal corresponding to a Cu(II) cluster. *a* denotes a *g*-value of 2.040 and corresponds to Cu(II). (**G**) Pchlide reduction activity of NB-protein and the BchN-BchB^H404A-M408A^ double mutant were assessed by adding L-protein and Pchlide (pre-bound to the respective NB-protein complex) in the presence of ATP-Mg^2+^. Alanine substitution of both His-404 and Met-408 residues that coordinate the Cu cluster completely abolishes the substrate reduction activity of DPOR.

Structures of *Rhodobacter capsulatus* NB-protein in the absence/presence of Pchlide have been solved using X-ray crystallography(*10, 12*). Similarly, a structure of the *Prochlorococcus marinus* NB-protein bound to L-protein in a 1:2 stoichiometry was trapped in the presence of ADP-AlF_3_ (**Fig. 1B**)(*7*). The general mechanistic assumption is that ET events in the two halves are uncoupled. However, we established that in both DPOR and nitrogenase, extensive allosteric communication exists between the two halves with events in one half regulating ET in the other(*13-15*). Recent Cryo-EM work on nitrogenase, performed under turnover conditions, show that the electron donor (Fe-protein) binds to only one half of the acceptor component complex (MoFe-protein), thereby setting asymmetry(*16*). In DPOR, we showed that the two halves of NB-protein function in an asymmetric manner initiated upon Pchlide binding(*17*). Thus, long-range allosteric communication between the NB-protein halves is required to coordinate asymmetry and to ensure proper delivery of the electron and to distinguish between substrate (Pchlide) and product (Chlide) that differ by a single double bond at the C17=C18 position (**Fig. 1A**). There are numerous steps in the DPOR catalytic cycle including Pchlide binding to both active sites in the NB-protein, engagement of ATP-bound L-protein to both halves of the NB-protein, ET, ATP hydrolysis, L-protein dissociation, and two repetitions of this cycle(*1, 11*). Here, we use Cryo-EM to capture the choreography of events and establish how the two halves function in an asymmetric manner.

We solved a 2.7 Å Cryo-EM structure of *Rhodobacter sphaeroides* NB-protein in the absence of Pchlide (**Figs. 1C, S1 & Table S1**). Overall, the arrangement of the BchN and BchB subunits are similar to the previously published *R. capsulatus* NB-protein structure with one [4Fe-4S]^NB^ cluster per half coordinated by 3 Cys and 1 Asp residues(*10*). For clarity, the following nomenclature is used to distinguish the BchN and BchB subunits in the two halves: BchN, BchB, BchN′, and BchB′. One major difference in our structure is the presence of strong additional density at the interface between the two halves (**Fig. 1D**). His-404 and Met-408 from both BchB and BchB′ protrude into this density and coordinate two Cu ions (**Fig. 1E**). The presence of Cu in the NB-protein was confirmed using inductively coupled plasma mass spectrometry (ICP-MS) (**Fig. S2**). The presence of Cu was further confirmed through EPR analysis of the NB-protein, and a signal was observed as a broad asymmetric line of ∼ 200 G width, centered around 3380 G, with absorption tailing off to the low field side of the line (**Fig. 1F**). The crossover field corresponds to a g-value of 2.040 and that value, along with the line shape, is interpreted as due to Cu(II). The large width of the line and lack of resolved ^63/65^Cu hyperfine structure is consistent with significant magnetic interaction, as expected in a binuclear cluster, and rules out a mononuclear copper source for the signal. In agreement with the EPR analysis, the additional density accommodates two Cu ions coordinated by the two His and Met residues along with two water molecules (**Fig. 1E**). To test whether the Cu cluster is important for substrate reduction, we substituted His-404 or Met-408 with Ala, or both in a double mutant (DPOR^H404A-M408A^), and measured Chlide formation in the presence of L-protein and ATP. The double mutant shows no presence of Cu in ICP-MS analysis (**Fig. S2**). While the single mutants are partially impaired for substrate reduction (**Fig. S3**), the double mutant is catalytically inactive (**Fig. 1G**) suggesting that the Cu cluster plays an essential role in DPOR activity.

To better resolve the role of the bi-copper cluster in substrate reduction, we solved a 3.1 Å Cryo-EM structure of the NB-protein bound to Pchlide (**Figs. 2A, S4 & Table S1**). Overall, the structure resembles the X-ray structure of Pchlide-bound NB-protein (PDB: 3AEQ & 3AEK) in terms of arrangement of the BchN and BchB subunits within the tetramer (*10*). However, there are several major differences: The first is observation of an intrinsic asymmetry between the BchN-BchB and BchN′-BchB′ halves (**Fig. 2B**). This is best visualized by comparing the overall differences in local resolution within the Cryo-EM maps (**Figs. 2B & C**). In the apo NB-protein structure, these changes are minimal (**Fig. 2C**); whereas in the Pchlide-bound NB-protein complex, such changes are more pronounced (dynamic) in one half (**Fig. 2B**). Interestingly, a path of dynamic changes can be traced from one half to the other and routes through the bi-copper cluster (**Fig. 2B**). This raises an intriguing possibility for establishment of an allosteric communication route between the two halves in DPOR upon substrate binding. The second observation identifies differences in the conformations between the Pchlide molecules bound in each active site (**Figs. 2D & E**). The porphyrin rings are bent in differing conformations suggesting that the active site likely probes and differentiates between substrate (Pchlide) versus product (Chlide) by sensing the differences in bendability. Since Pchlide and Chlide only differ by a single double-bond, local changes in energetics within the active site through bending/puckering likely drives binding of Pchlide and dissociation of Chlide. The structural asymmetry observed in the NB-protein upon binding of Pchlide also suggests that asymmetry is introduced or enhanced upon substrate binding in agreement with our previous biochemical observations(*17*).

**Figure 2.**
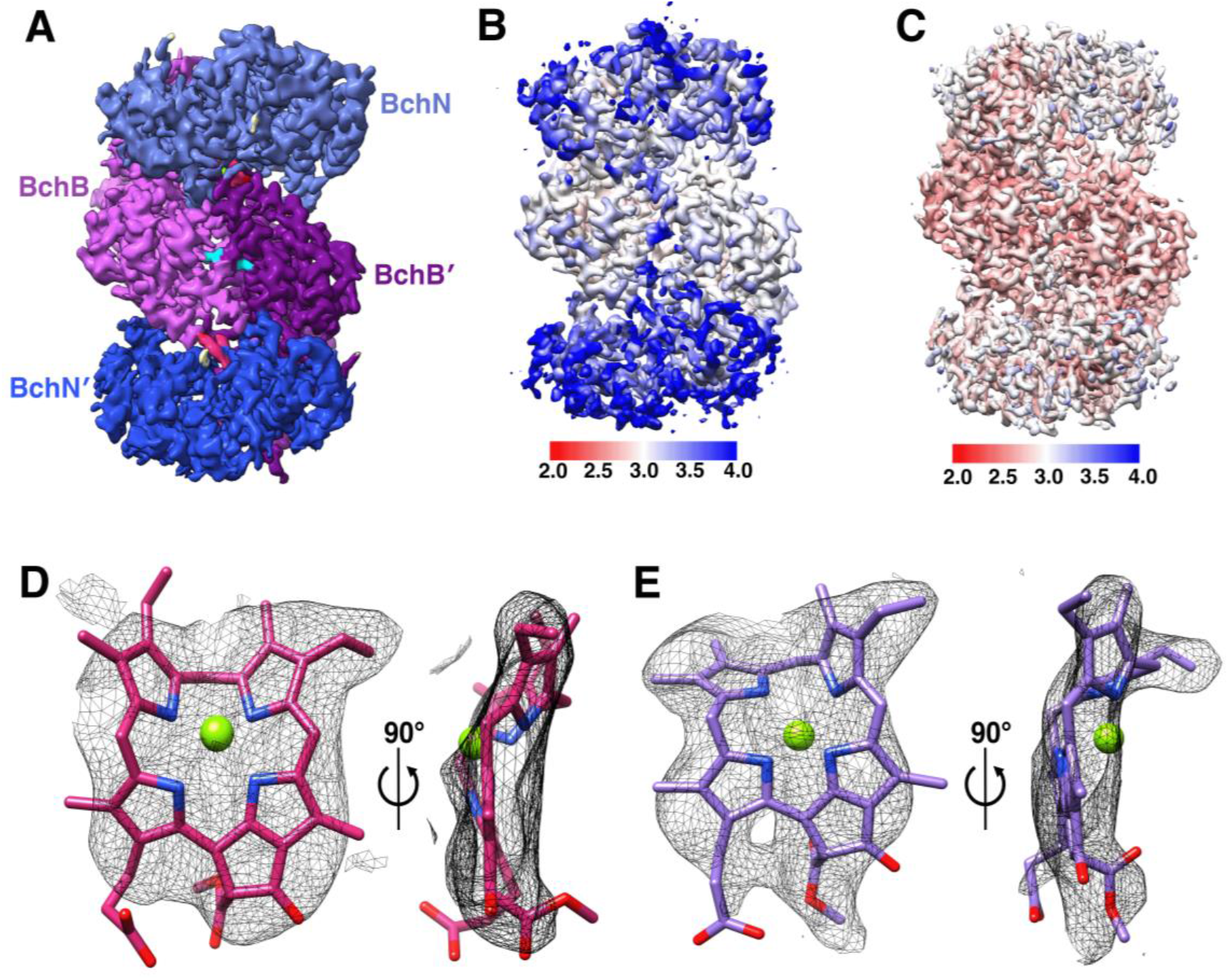
Pchlide binding enhances the asymmetry in the NB-protein and substrate recognition is achieved through puckering of the porphyrin ring. (**A**) A 3.1Å Cryo-EM map of *R. sphaeroides* NB-protein bound to Pchlide. (**B**) Local resolution analysis of the NB-protein-Pchlide complex is denoted as a heatmap where red and blue denote regions of higher-resolution (less-dynamic) versus lower-resolution (dynamic), respectively. Asymmetry in the NB-protein is visible through more dynamic changes in one half of the complex. An intrinsic path of dynamic changes can be traced from one half of the NB-protein to the other displaying a potential path of communication that traverses through the bi-copper cluster. (**C**) Local resolution map for the apo NB-protein displays lesser dynamic changes compared to the Pchlide-bound NB-protein complex in panel B. The top half is slightly more dynamic compared to the lower half of the NB-protein suggesting that asymmetry might be an inherent property that is enhanced upon Pchlide binding. (**D & E**) Carved electron density for the Pchlide molecules bound to each active site in the NB-protein. The Pchlide molecules are non-planar and show puckering of the porphyrin rings. The electron densities for the two Pchlide molecules reveal different conformations in each half suggesting asymmetric substrate binding/recognition properties. The Pchlide molecules are colored pink and purple to denote the two active sites in the NB-protein tetramer.

The bi-copper cluster is well ordered in the Pchlide-bound NB-protein complex. In the apo structure, Met-408 adopts two conformations, but becomes ordered upon Pchlide binding (**Fig. S5)**. His-404 and Met-408 reside in a long α-helix (residues 391-418) in BchB. This helix promotes interaction in *trans* where residues from BchB from one half interacts with Pchlide bound in the active site of the other BchNB′ half (**Fig. S6**). In the previously solved X-ray structures of NB-protein, there are no reports of bound Cu (**Figs. S6A-C**)(*7, 10, 12*). For some of these structures, NB-proteins were endogenously purified from *Rhodobacter capsulatus* (*10*). Thus, we reassessed the deposited electron density maps of NB-protein bound to Pchlide (PDB: 3AEQ/3AEK) and indeed find additional density at the tetramer interface. As deposited, water molecules are modeled into the additional density (**Figs. S6B**). We can model two Cu molecules within this density with excellent coordination by two His residues (**Fig. S6D**). Surprisingly, in the apo structure (PDB:3AER), this additional density is missing (**Fig. S6A**). Crystallization conditions for the apo structure contained 0.2M ammonium chloride(*10*) and ammonium salts are known to interact with and precipitate Cu(*18*). Thus, over the course of crystallization, Cu was likely sequestered away by ammonium salts. Knowledge of the functional role(s) of Cu is extremely important because of the nature of the two helices on either side of this cluster (**Figs. S6**). In the X-ray structures, this helix is unbent in the apo structure, but is bent in the Pchlide bound structure (**Fig. S6I**). In contrast, this helix is intact and does not bend in our Cryo-EM structures (**Figs. S6J**). The bending of this helix has been proposed as a mechanism for substrate binding in *R. capsulatus* DPOR(*10*). However, we observe no evidence of such bending in *R. sphaeroides* DPOR. Met-404 and His-408 are well conserved in *Rhodobacter* (**Fig. S7**), thus how this helix mediates substrate binding and reduction remains to be established.

To better understand the role of asymmetry in the NB-protein, the bi-copper cluster, and distortion of the helix (discussed above) in substrate reduction, we solved a 3.7 Å Cryo-EM structure of DPOR under turnover conditions (**Figs. 3A, S8 & Table S2**). Here, Pchlide-bound NB-protein was mixed with L-protein and ATP-Mg^2+^ for 5 minutes, applied onto the grids, frozen and imaged. The 2D classification analysis of this dataset shows unbound L-protein and NB-protein molecules and NB-protein tetramers bound to one L-protein dimer (**Figs. 3A & S8**). However, we observe no evidence for the hetero-octameric (BchL)_2_-(BchN-BchB)_2_-(BchL)_2_ complex as captured for *Prochlorococcus marinus* DPOR in the presence of ADP-AlF_3_ (PDB: 2YNM; **Fig. 1B**)(*7*). Our structure shows that L-protein binding to NB-protein during turnover is asymmetric and this finding is similar to recent observations for nitrogenase during turnover where only one Fe-protein is bound to the MoFe-protein(*16*). To test whether symmetric BchL binding is a structural feature driven by ADP-AlF_3_, we solved a 3.7 Å Cryo-EM structure of *R. sphaeroides* DPOR in the presence of ADP-AlF_3_ (**Figs. 3B, S9 & Table S2**). Under these conditions, 2D classification shows free L-protein and NB-protein molecules in addition to two L-protein bound NB-protein complexes (**Fig. S9**). Surprisingly, we see no evidence for the one L-protein bound NB-protein complexes in 2D classification. To understand the differences in complex formation during turnover (ATP-Mg^2+^) versus in the presence of ADP-AlF_3_, we compared the differences in the local resolution within the two structures (**Figs. 3C & D**). In the turnover data, asymmetric dynamic changes are pronounced in one half compared to the other and L-protein binding induces relative stability to one half of the NB-protein. Such asymmetric changes are predominantly lost in the presence of ADP-AlF_3_. Nevertheless, the path of allosteric changes is strikingly visible in both structures and route through the bi-copper cluster at the NB-protein interface. In addition, the conformation of the Pchlide molecules bound to the two active sites in the turnover versus transition complexes show striking differences (**Figs. 3E-H**). In the turnover complex, the two Pchlide molecules are puckered with distinct conformations (**Figs. 3E & F**). In contrast, in the transition-state complex, while the two Pchlide molecules show differences, they are puckered similarly (**Figs. 3G & H**). We also note that the Pchlide molecules appear to be more dynamic in the transition-state DPOR complex as additional unresolved density is present adjacent to the active site in both halves of the NB-protein (**Fig. S10**). This suggests that asymmetric control of substrate binding, recognition, and electron transfer are lost when the DPOR complex is assembled in the presence of ADP-AlF_3_.

**Figure 3.**
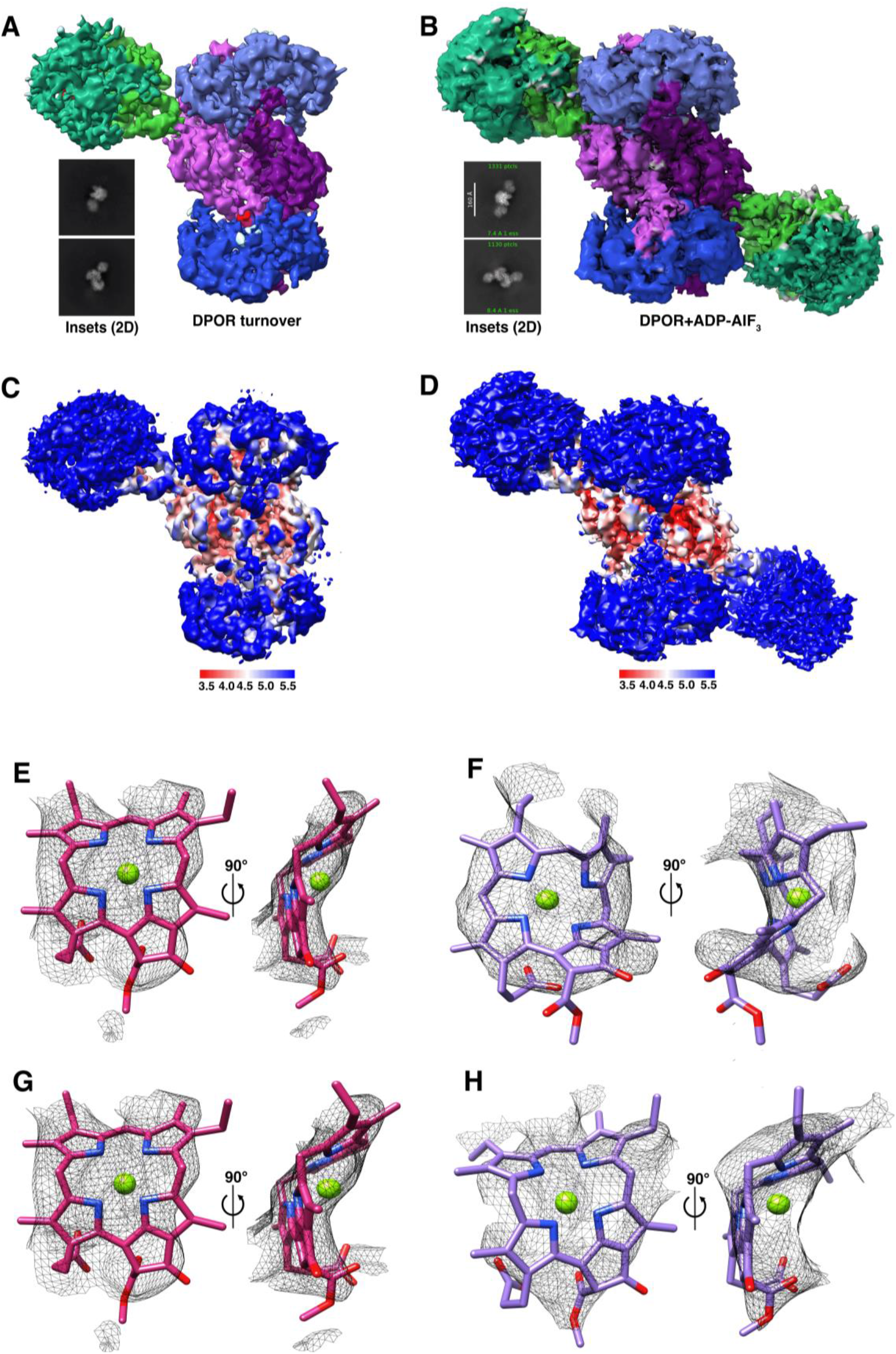
Cryo-EM map of DPOR under turnover shows asymmetric binding of L-protein to the NB-protein. (**A**) A 3.7Å Cryo-EM map of DPOR solved under turnover conditions in the presence of ATP is shown along with representative 2D classes. During turnover, at 5 minutes post addition of ATP, only one L-protein is bound to the NB-protein. The two BchL subunits (L_1_ and L_2_) are shown in dark and light green, respectively. (**B**) A 3.7 Å Cryo-EM map of DPOR in the presence of ADP-AlF_3_ is shown along with representative 2D classes. In this transition-state complex, two L-proteins are bound to the NB-protein. Local resolution maps for the structures of DPOR under (**C**) turnover and (**D**) transition-state are shown. Both structures show robust dynamic changes. Stark asymmetric dynamic changes are observed in the turnover complex. The asymmetry, while present, is considerably muted in the transition-state complex. Under both conditions, the path of allosteric communication between the two halves is prominently observable and routes through the bi-copper cluster in the NB-protein. Structure of the Pchlide molecules bound in the **E)** BchN-BchB or **F)** BchN’-BchB’ active sites in the DPOR-turnover CryoEM structure. The conformations of the two substrate molecules are different suggesting asymmetric interactions. Structure of the Pchlide molecules in the **G)** BchN-BchB or **H)** BchN’-BchB’ active sites in the DPOR-transition-state CryoEM structure shows largely similar puckering of the two substrate molecules suggesting that asymmetry is predominantly suppressed. In both structures, clear evidence for puckering of Pchlide is observed suggesting a potential mechanism of differentiation between substrate (Pchlide) and product (Chlide).

In both our turnover and ADP-AlF_3_ DPOR structures, the position of L-protein bound on NB-protein is rotated by ∼45-50° in comparison to the crystal structures (**Figs. 4A & S11**). Thus, the distance between the [4Fe-4S] clusters between L-protein and NB-protein is ∼39Å compared to ∼14Å in the crystal structure (PDB: 2YNM; **Fig. S12**). Closer positioning of the two FeS clusters likely promotes ET. Thus, a rolling motion of L-protein between the two states would be required for ET, as established for the nitrogenase complex(*13, 16, 19-22*). We had previously uncovered that a disordered region in the N-terminus of L-protein is autoinhibitory to function and binds across the [4Fe-4S]^L^ cluster(*23*). ATP binding to L-protein remodels this interaction and promotes L-protein binding and subsequent ET to the NB-protein. In the Cryo-EM structures, the two BchL subunits are positioned such that they make contact with only the BchB subunit with the two disordered regions crossing over to regulate protein-protein interactions (**Figs. 4B & C**). In the turnover complex, the binding between L-protein and NB-protein is promoted through several salt-bridges between L-protein and the BchB subunit (**Fig. 4B**). In the ADP-AlF_3_ trapped complex, additional contacts further stabilize the binding, suggesting a slightly different binding conformation (**Fig. 4C**). Interestingly, these contacts vary between the two docked L-proteins in contrast to symmetrical contacts observed in the X-ray structure (PDB:2YNM)(*7*).

**Figure 4.**
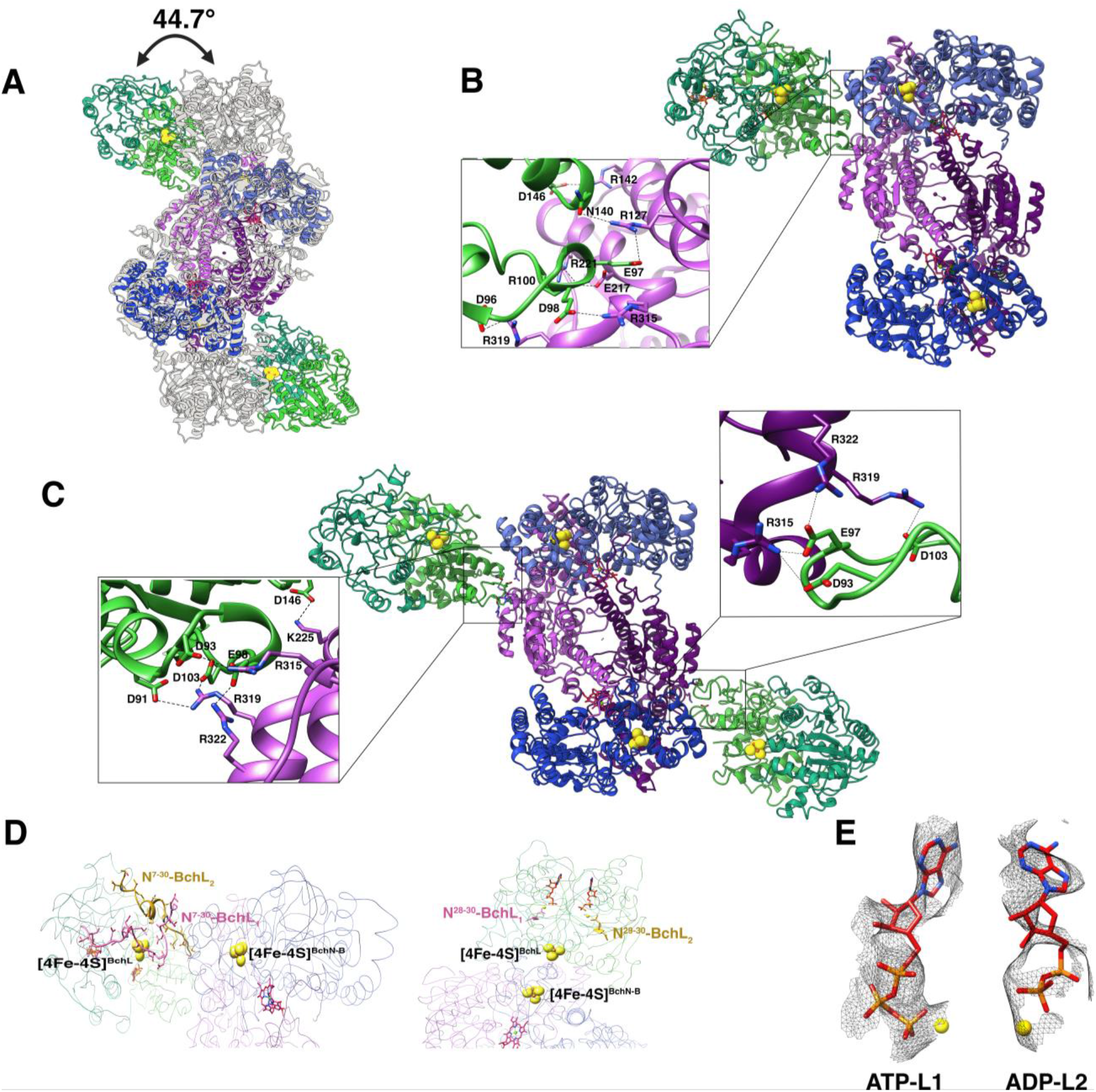
Rolling-motion of the L-protein on the NB-protein is regulated by the disordered N-terminal region and ATP binding/hydrolysis. (**A**) Superposition of the *P. marinus* transition-state DPOR crystal structure (PDB: 2YNM in grey) with the Cryo-EM structure (both solved in the presence of ADP-AlF_3_) shows a large 45º difference in the docking positions of the respective L-proteins on the NB-protein. Salt-bridge interactions between the L-protein and the NB protein in the (**B**) turnover complex and (**C**) transition-state complex are shown. (**D**) In the Cryo-EM structure of DPOR under turnover conditions (left), the disordered N-terminal regions of BchL (colored yellow; residues 7 to 30) are observed and situated at the binding interface between the two component proteins. This region is not observed in the crystal structure of the *P. marinus* DPOR complex (PDB:2YNM, right). (**D**) Carved electron density for ATP bound to the L_1_ subunit of L-protein versus the ADP bound to L_2_.

We had previously shown that a disordered N-terminal region in BchL serves a inhibitory role by binding across the [4Fe-4S]^L^ cluster and likely preventing complex formation with the NB-protein(*23*). We had established that ATP binding to the L-protein remodels this interaction and leading to complex formation with the NB-protein. However, the functional significance of this feature was poorly understood. In our Cryo-EM structures, the [4Fe-4S]^L^ cluster is buried by the disordered regions (**Fig. 4D**). This inhibition/protection of the cluster likely explains the slow kinetics of Pchlide reduction by DPOR (**Fig. 2G**). Since the L-protein is dynamic in our Cryo-EM structure, precise determination of ATP versus ADP bound during catalysis is difficult. However, comparison of the densities within the two active sites in the L-protein suggests asymmetic states of nucleotide hydrolysis. Clear density for the three phosphates of ATP is observed in one subunit, but density for the third phosphate is missing in the other (**Fig. 4E**). Thus, binding and rolling of L-protein on the NB-protein is coupled to asymmetric ATP binding and hydrolysis within the L-protein and further regulated by the disordered N-terminal region.

We observe introduction of asymmetry in the NB-protein upon Pchlide binding and further propagated by asymmetric binding of L-protein. Thus, we sought to understand how asymmetry is introduced, maintained, and used for ET and substrate reduction in DPOR. Aromatic residues are commonly used as relays for long-range ET(*24*). Thus, we analyzed the conformational changes in key aromatic residues in the NB-protein situated between the [4Fe-4S] clusters and the bi-copper center (**Fig. 5A**). An overlay of the structural changes in NB-protein between the apo, Pchlide-bound, and turnover Cryo-EM structures reveals a snapshot of how allosteric communication is likely propagated in DPOR. A direct comparison of these changes between the two halves reveals asymmetric movements in key residues (**Supplemental Video 1**). Trp-6, Phe-28, Trp-36, Phe-67, Tyr-277, Trp-290, Phe-396, and Phe-418 all show major asymmetric sidechain conformations (**Fig. 5A**). In addition, the imidazolium ring of His residues also participate in ET(*25*). His-13, His-378, and His-56 residues all display asymmetric side-chain conformations (**Fig. 5A**). Finally, His-404 and Met-408 that coordinate the bi-copper cluster also display asymmetric conformational changes suggesting that the Cu ions and the associated helix are integral to allosteric control in DPOR (**Supplemental Video 1**). How the redox state of the bi-copper cluster serves as a relay switch to regulate/communicate allostery across the two subunits remains to be established.

**Figure 5.**
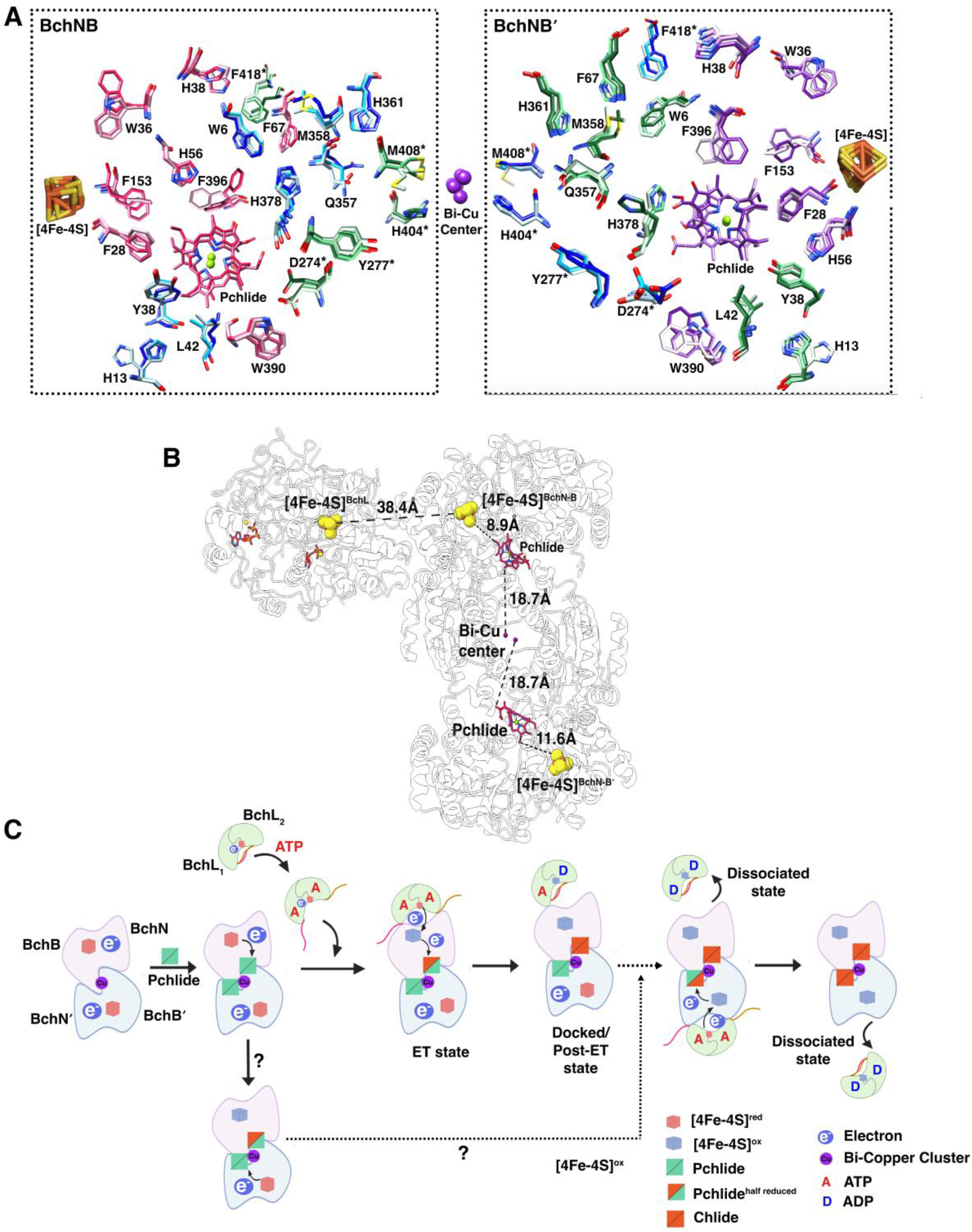
A proposed mechanism for the role of asymmetry in electron transfer. (**A**) A superposition of the NB-protein from the three Cryo-EM structures (apo, Pchlide-bound, and under turnover) is shown and the key residues, the [4Fe-4S]^NB^ cluster, and the bi-copper cluster are depicted. Overall changes in the environment surrounding the substrate and clusters are visible, and asymmetric motions are observed between the two halves of the NB-protein. The residues are colored to match the individual NB-protein subunits and shaded to correspond the NB-protein apo (light), Pchlide-bound NB-protein (medium), and DPOR turnover (dark) structures. Residues colored pink, blue, purple, and green represent BchN, BchB, BchN′, and BchB′, respectively. The * denotes residues that interact across the two halves *in trans*. We propose that transfer of an electron in the left half of the NB-protein (bound to L-protein) is promoted due to proper alignment interactions that favor ET. These residues are not aligned in the right half leading to the allosteric suppression of L-protein binding and conditions that do not favor ET. (**B**) The relative distances between the [4Fe-4S] clusters, Pchlide molecules, and the bi-copper center is denoted on the DPOR structure under turnover conditions. A ∼2-3 Å difference between the [4Fe-4S]^NB^ cluster and the Pchlide within the two halves in the NB-protein is observed. (**C**) A proposed model for asymmetric events in the ET cycle in DPOR is shown. Release of the disordered N-terminal regions in the L-protein upon ATP binding is a prerequisite for binding to the NB-protein to initiate ET. Since the [4Fe-4S]^NB^ cluster is pre-reduced, the first electron is transferred to Pchlide through a deficit-spending mechanism. Whether this happens on both sides, or allosterically suppressed in one half, remains to be elucidated. Binding of reduced L-protein drives the second ET event leading to Pchlide reduction in one active site. The timing of L-protein binding to the other half and the associated regulatory events are to be uncovered.

DPOR and other nitrogenase-like enzymes share common overall structural resemblance to nitrogenase. Recent Cryo-EM data on nitrogenase along with data for DPOR presented here show that these protein complexes function asymmetrically(*16, 26*). For DPOR, asymmetry is introduced upon Pchlide binding to BchNB which then dictates which half of the NB-protein interacts with L-protein. Based on our structural work, we propose that allosteric communication can transpire through dual paths. The first arises from changes within the active site upon Pchlide binding that is then propagated to the L-protein binding interface on NB-protein. The second level of regulation is initiated upon L-protein binding which likely suppressess L-protein binding on the opposite BchNB′ interface which is situated ∼100Å away. The discovery of the bi-copper cluster at the middle of the NB-protein tetramer and the associated helices that interact with Pchlide and the active sites in *trans* adds another layer of intrigue to the ET and substrate reduction mechanisms. The relationships between electron flow from the [4Fe-4S]^NB^ cluster of the NB-protein to Pchlide and/or from the bi-copper cluster remains to be resolved. The Pchlide is situated ∼18.7Å from the bi-copper cluster (**Fig. 5B**) compared to ∼9-12Å from the [4Fe-4S]^NB^ cluster of NB-protein. The [4Fe-4S]^NB^ cluster of NB-protein is pre-reduced and we have shown that this electron is transferred to Pchlide in the absence of L-protein using a deficit spending like mechanism shown for nitrogenase(*17*). This is also evident in our single-turnover Pchlide reduction experiments where half the Pchlide is reduced rapidly (within 10 minutes) wheres the reduction of the second Pchlide takes more than an hour (**Fig. 1G**). This would suggest that the enzyme is able to count the electrons transferred through conformational changes and/or bending/puckering of the Pchlide molecule. While speculative, we envision that transfer of the first electron upon Pchlide binding induces the asymmetry and differential bending of the Pchlide molecules we observe in the Cryo-EM structures (**Figs. 2 & 3**). The second electron required to complete Pchlide reduction is provided upon L-protein binding. This second ET step transpires rapidly in one half. The long-range allosteric changes and negative cooperativity occur on a much slower time scale resulting in slower binding of L-protein on BchNB′. Another puzzle is the coupling of ATP utilization to the remodeling of the inhibitory N-terminal region of the L-protein. We propose that ATP binding to the two BchL subunits probably releases the inhibition in both subunits resulting in the rapid docking and immediate ET (**Fig. 5C**). The subsequent asymmetric ATP hydrolysis in one BchL subunit triggers the rolling motion to a post-ET state where the N-terminal disordered region has coallapsed to the inhibitory state and is captured in our Cryo-EM structures. When and how this movement triggers L-protein binding in the opposite half of BchNB′, along with a host of other mechanistic queries, remain a fascinating mystery to be interrogated: How does the bi-copper cluster and its redox state control asymmetry? How and when is ET activated in the suppressed half? Does this occur upon ATP hydrolysis and L-protein rolling/dissociation in the first BchNB half upon Pchlide to Chlide reduction? The structural work presented here opens up a fascinating series of mechanistic questions regarding the mechanisms of allosteric control in long-range ET in nitrogenase-like enzymes.

## Acknowledgments

We thank Dr. Summer Brock and Dr. Katherine Basore at the Washington University Center for Cellular Imaging and Dr. Liguo Wang at the Laboratory for BioMolecular Structure (LBMS) at Brookhaven National Labs for additional Cryo-EM data collection. We also thank Dr. Bassem Mohammed for advice on data processing.

## Funding

Department of Energy, Office of Basic Energy Sciences, DE-SC0020965 (EA)

Department of Energy, Office of Basic Energy Sciences, KP1607011 (LBMS, BNL)

National Science Foundation, CHE 2204023 (BB)

National Institutes of Health, Office of the Director, S10 OD030343 (EA)

## Author contributions

Conceptualization: EA, RK

Methodology: RK, JD, MTL, BB, EA

Investigation: RK, NW, JD, NW, BB, EA

Visualization: RK, JD, EA

Funding acquisition: EA, BB, BB

Project administration: EA

Supervision: EA, RK, BB

Writing – original draft: EA, RK

Writing – review & editing: EA, RK, JD, NW, BB, BB

## Data and materials availability

NB-protein apo **PDB: 8VQH & EMDB-43443**, Pchlide-bound NB-protein **PDB: 8VQI & EMDB-43444**, DPOR under turnover **PDB: 8VQJ & EMDB-43446**

## Supplementary Materials

Materials and Methods

Figs. S1 to S12

Tables S1 to S2

Movies S1 References (*1*–*11*)

