## Supplemental Information for "Cryo-EM captures the coordination of long-range allostery and asymmetric electron transfer by a bi-copper cluster in the nitrogenase-like DPOR complex"

### Materials and Methods

*Plasmids:* Plasmids for recombinant overproduction of BchL and BchNB were previously described (*1-4*). Briefly, the open reading frames for BchB and BchN were PCR amplified from *Rhodobacter sphaeroides* genomic DNA and cloned into the multiple cloning sites 1 and 2 of a pRSF-Duet1 vector, respectively. BchB was designed to carry an N-terminal 6X poly-histidine tag followed by a 3C protease cleavage site. Please note that addition of an affinity tag to either the N- or C-terminus of BchN will lead to loss in activity (*2*). The BchL open reading frame was similarly PCR amplified and cloned into pRSF-Duet1 with an N-terminal 6X poly-histidine tag followed by a 3C protease cleavage site. Point mutants described were generated using Q5 site-directed mutagenesis (New England Biolabs).

*Protein Expression and Purification:* Wild-type and mutant DPOR component proteins (L-protein and NB-protein) were purified separately using *SufFeScient E. coli* BL21(DE3) cells (PK11466) as previously described (*1-4*) with minor modifications. Transformants were grown in 4.5 L Luria-Bertani (LB) medium, supplemented with 1 mM of L-cysteine and 1 mM of iron (III) citrate. Cultures were grown at 37 °C with shaking until the optical density of the cultures at 600 nm reached 0.6-0.8. Protein expression was induced by the addition of Isopropyl β-D-1-thiogalactopyranoside (IPTG) to a final concentration of 0.1 mM. Cultures were grown for an additional 12 h with shaking at 17 °C. The cells were transferred to centrifuge bottles with rubber gaskets and incubated with 1.7 mM sodium dithionite and 1 mM $CuCl_2$ for 3 h at 17 °C. Cells were harvested by centrifugation, and the bottles were moved into a Coy Lab anaerobic chamber (glove box) under a 95% $N_2$/5% $H_2$ mixture before decatenation. All further lysis and purification steps were performed under anoxygenic conditions and inside the glove box as previously described (*1-3, 5*). All buffers were purged of air under ultrahigh-purity $N_2$ on a home-built Schlenk line and stored under $N_2$. Cell pellets were resuspended in ~100 ml of Buffer A (100 mM HEPES pH 7.5, 150 mM NaCl, 10 mM $MgCl_2$, 10% glycerol, and 1.7 mM of sodium dithionite) supplemented with 3X protease inhibitor cocktail (Sigma Inc) and 0.1 mg/mL Lysozyme. Cells were lysed by sonication (pulse of 1 sec ON and 3 sec OFF for a total of 3 min) and the lysate was centrifuged at 37,157xg for 60 min at 4 °C. The clarified supernatant was loaded on to a 5 mL $Ni^{2+}$-NTA column (Gold Biotechnology Inc.) equilibrated with Buffer A. Both NB-protein and L-protein complexes were eluted with a step-gradient of imidazole. Fractions containing protein were brown in color and were analyzed on SDS-PAGE for purity (>95% pure). These fractions were pooled and concentrated using an Amicon spin concentrator (30 kDa cutoff) and further fractionated on S200 HiLoad 26/600 size exclusion chromatography column (GE healthcare) pre-equilibrated with Buffer A containing no glycerol. Protein containing fractions were pooled, concentrated, and aliquoted into 1.2 ml cryo-tubes with a gasket sealed cap (Corning Scientific). Closed tubes with protein were flash frozen in the glove box gas exchanger chamber using liquid nitrogen and stored under liquid nitrogen. Protein concentrations were determined using Bradford reagent with BSA as the reference.

*Generation of Pchlide:* Pchlide was generated from a *Rhodobacter capsulatus* ZY-5 strain harboring a deletion of the BchL gene (*6, 7*) (a kind gift from Dr. Carl Bauer, Indiana University) and purified as described(*1-4*).

*Preparation of Pchlide-bound NB-protein for Cryo-EM and substrate reduction analysis:* NB-protein (67 μM tetramer; 33.5 μM substrate binding sites) was mixed with a slight excess of Pchlide (40 μM) and passed over a SEC column (Superose 6 Increase 5/150 GL; Cytiva Inc.). Peak fractions containing Pchlide-bound NB-protein were were pooled and concentrated to ~20 μM using an Amicon spin concentrator (30 kDa cutoff). Protein concentration was estimated using a Bradford assay with BSA as reference.

*Pchlide reduction assay*: 20 μM NB-protein (tetramer concentration; 40 μM total L-protein binding sites) pre-bound to Pchlide (as described above) was mixed with 80 μM L-protein (dimer concentration) in buffer A (100 mM HEPES pH 7.5, 150 mM NaCl, 10 mM $MgCl_2$, 10% glycerol, and 1.7 mM of sodium dithionite). 1 ml of total reaction was made, and reactions initiated by the addition of 3 mM ATP (final concentration). At the defined time points, 200 μl of the reaction was extracted and quenched with 800 μl of 100% acetone. The acetone/reaction mixture was spun down in a table-top centrifuge at 13,226xg for 4 min. The supernatant was transferred to a cuvette and absorbance scans (600 nm to 700 nm) were recorded on a Cary 100 UV-Vis spectrophotometer using quartz cuvettes (Agilent Technologies). Chlide formation was quantified using molar extinction coefficient $\varepsilon_{666nm}$=74,900 $M^{-1}cm^{-1}$. The Pchlide reduction traces were normalized using background spectra recorded in the absence of ATP.

*ICP-MS:* The metal analysis was performed on 7800 ICP-MS (Agilent Technologies) coupled to an SPS4 autosampler (Agilent Technologies) as previously described (*8*). Briefly, the wild-type and mutant DPOR complexes were diluted to the same concentration and 1435 µg of each complex was digested in 5% optima grade $HNO_3$ (Fisher Chemical) at 99 °C for 30 min. After incubation, samples were cooled to room temperature, briefly spun down, and 100 µL (around 1400 µg of protein) of the clear supernatant was diluted to 4 mL (2% HNO3, 0.5% HCl in water) for metal analysis. The signal for $^{63}Cu$ ions were monitored for 30 sec with an integration time/mass of 1.5 sec/ion in helium mode. The mobile phase was 2% $HNO_3$, 0.5% HCl in water, with a flow rate of 1 mL/min. The 4 mL volume allowed for collection of two measurements per complex (n = 2).

*Sample Preparation for Cryo-EM:* For the turnover complex, Pchlide-bound NB-protein and L-protein were mixed as described above for the Pchlide reduction assay reactions, but in a smaller total volume (10 µl). The sample was prepared in a sealed glass vial inside the anaerobic chamber. Similarly, a stock of 100 mM ATP in Buffer A was prepared in a separate sealed glass vial. ATP was mixed with the DPOR protein using a Hamilton gas tight syringe and incubated for 5 minutes and immediately applied onto the grid. The NB-protein apo, Pchlide-bound, or the transition-state (ADP-$AlF_3$) samples were also processed similarly under anaerobic conditions, but without the need for separate mixing of nucleotide. Cryo-EM samples were prepared on Quantifoil R2/2 (Q350CR2, Electron Microscopy Sciences) holey carbon grids that were plasma cleaned for 60 s using a Solarus 950 (Gatan) with an $H_2/O_2$ mixture. Samples were applied onto the grids using a

Vitrobot Mark IV (Thermo Fisher Scientific) with the chamber at 4°C and 100% humidity. To reduce the exposure to the air, a Hamilton gas tight syringe was used to rapidly apply 3 µl of the sample on the grid from the sealed glass vial. After a 20 s incubation, each grid was blotted for 2 s with a blot force of 0 and immediately plunged into liquid ethane cooled by liquid nitrogen. The total time that each sample spent outside of sealed glass vial during grid preparation and freezing was less than 50 sec. Grids were stored under liquid $N_2$ until data collection. Grids were clipped and loaded into an image corrected 300 kV Titan Krios G3i Cryo-TEM (Thermo Fisher Scientific). Movies were recorded with a Gatan K3 Bioquantum direct electron detector (Gatan Ametek) in an automated fashion using EPU software 2.12.1 (Thermo Fisher Scientific) with a pixel size of 0.825 Å and a nominal defocus range of -1.0 to -2.4 µm. Data were acquired with a dose rate of 22 $e^-/Å^2$ over a 2.3 s exposure time, yielding a total dose of 50 $e^-/Å^2$ over 40 fractions. This sample preparation strategy was optimal in achieving consistent uniform thin ice and no air-water interface issues were encountered. Excellent distribution of diverse and alternate views of the protein in 2D classification were observed for all the samples.

*Image processing:*
**NB-protein apo structure**: The single-particle cryo-EM data was processed using cryoSPARC v4.4.1. For the NB-protein apo dataset, 4376 raw movies were motion corrected using patch motion correction and CTF estimation was performed using patch CTF estimation. Particles were picked initially using blob particle picking and extracted using a box size of 400 pixels. The resulting particle stack (2,532,257 particles) was subjected to several rounds of 2D classification to generate an initial clean particle stack (551,140 particle) for 3D construction using *ab initio*. Initial 3D volumes were further cleaned using heterogeneous refinement. Volumes were examined in ChimeraX (*4*) and a 3D volume-particle stack combination (357,713 particles) was selected for further processing. Low quality particles in the stack were further removed using several rounds of 2D classification. This was followed by non-uniform refinement of the consensus volume using 160,048 particles (final particle stack) with well-defined NB-protein density. The resulting cryo-EM density yielded a map with an overall resolution of 2.7 Å at a Fourier shell correlation cut-off of 0.143.

**NB-protein bound to Pchlide**: For the Pchlide-bound NB-protein dataset, 5394 raw movies were motion corrected using patch motion correction and CTF estimation was performed using patch CTF estimation. Particles were picked initially using blob particle picking and extracted using a box size of 400 pixels. The resulting particles stack (3,107,286 particles) was subjected to several rounds of 2D classification to generate an initial clean particle stack (342,672 particle) for initial 3D construction using *ab initio*. Initial 3D volumes were further cleaned using heterogeneous refinement. Volumes were examined in ChimeraX (*4*) and a 3D volume-particle stack combination (160,124 particles) was selected for further processing. Low quality particles in the stack were further removed using several rounds of 2D classification. This was followed by non-uniform refinement of the consensus volume using 72,187 particles (final particle stack) with well-defined NB-protein and bound-Pchlide density. The resulting cryo-EM density yielded a map with an overall resolution of 3.02 Å at a Fourier shell correlation cut-off of 0.143.

**DPOR structure under turnover**: For the DPOR-turnover (ATP-$Mg^{2+}$) dataset, 3725 raw movies were motion corrected using patch motion correction and CTF estimation was performed using patch CTF estimation. Particles were picked initially using template picker with the NB-protein structure as an input model and extracted using a box size of 480 pixels. The resulting particle stack (856,671 particles) was subjected to several rounds of 2D classification to generate an initial clean particle stack (190,994 particle) for 3D construction using *ab initio*. Note that the initial 2D classes were a combination of unbound L-protein and NB-protein along with a complex of NB-protein and L-protein at a 1:1 ratio (a NB-protein tetramer bound to one L-protein homodimer). Surprisingly, no 2D classes showing two L-protein homodimers bound to the NB-protein tetramer (as seen in former X-ray crystallography studies) were observed. 2D classes corresponding to the complex were selected and processed further. Initial 3D volumes were further cleaned using heterogeneous refinement. Volumes were examined in ChimeraX (*4*) and a 3D volume particle stack combination (84,350 particles) was selected for further processing. This was followed by homo-refinement with a particle stack of 83,885 particles. The particle stack was further 3D classified into four classes followed by hetero refinement on a particle stack of 21,290 particles. This was followed by final homo refinement of the consensus volume using 18,965 particles (final particle stack) with well-defined NB-protein with Pchlide and bound to only one L-protein homodimer. The resulting cryo-EM density yielded a map with an overall resolution of 3.82 Å at a Fourier shell correlation cut-off of 0.143.

**DPOR structure under transition state**: For the DPOR dataset collected under turnover conditions (ADP-$AlF_3$), 10,488 raw movies were motion corrected using patch motion correction and CTF estimation was performed using patch CTF estimation. Particles were picked initially using template picker with a model of DPOR as input and extracted using a box size of 480 pixels. The resulting particle stack (1,071,772 particles) was subjected to several rounds of 2D classification to generate an initial clean particle stack (73,354 particle) for initial 3D construction using *ab initio*. Note that the initial 2D classes were a combination of unbound L-protein and NB-protein along with a complex of NB-protein and L-protein at a 1:2 ratio (a NB-protein tetramer bound to two L-protein homodimers). In contrast to the turnover dataset, no 1:1 complexes were observed. 2D classes of the 1:2 complex were selected for further processing. Initial 3D volumes were further cleaned using heterogeneous refinement. This was followed up by homo- and non-uniform refinement with a particle stack of 44,953 particles. The resulting particle stack was further 3D classified into two classes followed by homo refinement on a particle stack. This was followed by final non-uniform refinement of the consensus volume using 20,571 particles (final particle stack) with well-defined NB-protein bound to Pchlide and two BchL dimers. The resulting cryo-EM density yielded a map with an overall resolution of 3.68 Å at a Fourier shell correlation cut-off of 0.143.

*Model building and refinement:* ChimeraX was used to dock the NB-protein (PDB: 3AER) model into the electron microscopy density maps for NB-protein apo and NB-protein Pchlide-bound complexes. Similarly, L-protein (PDB:6UYK) along with NB-protein (PDB: 3AER) models were combined and docked in the electron density for the full DPOR complex under turnover and transition-state conditions. Subsequent models were iteratively refined and rebuilt in COOT-0.9.8.5 EL(*9*) and further refined and validated using Phenix. Structural figures were generated using Chimera(*10*) or ChimeraX(*11*).

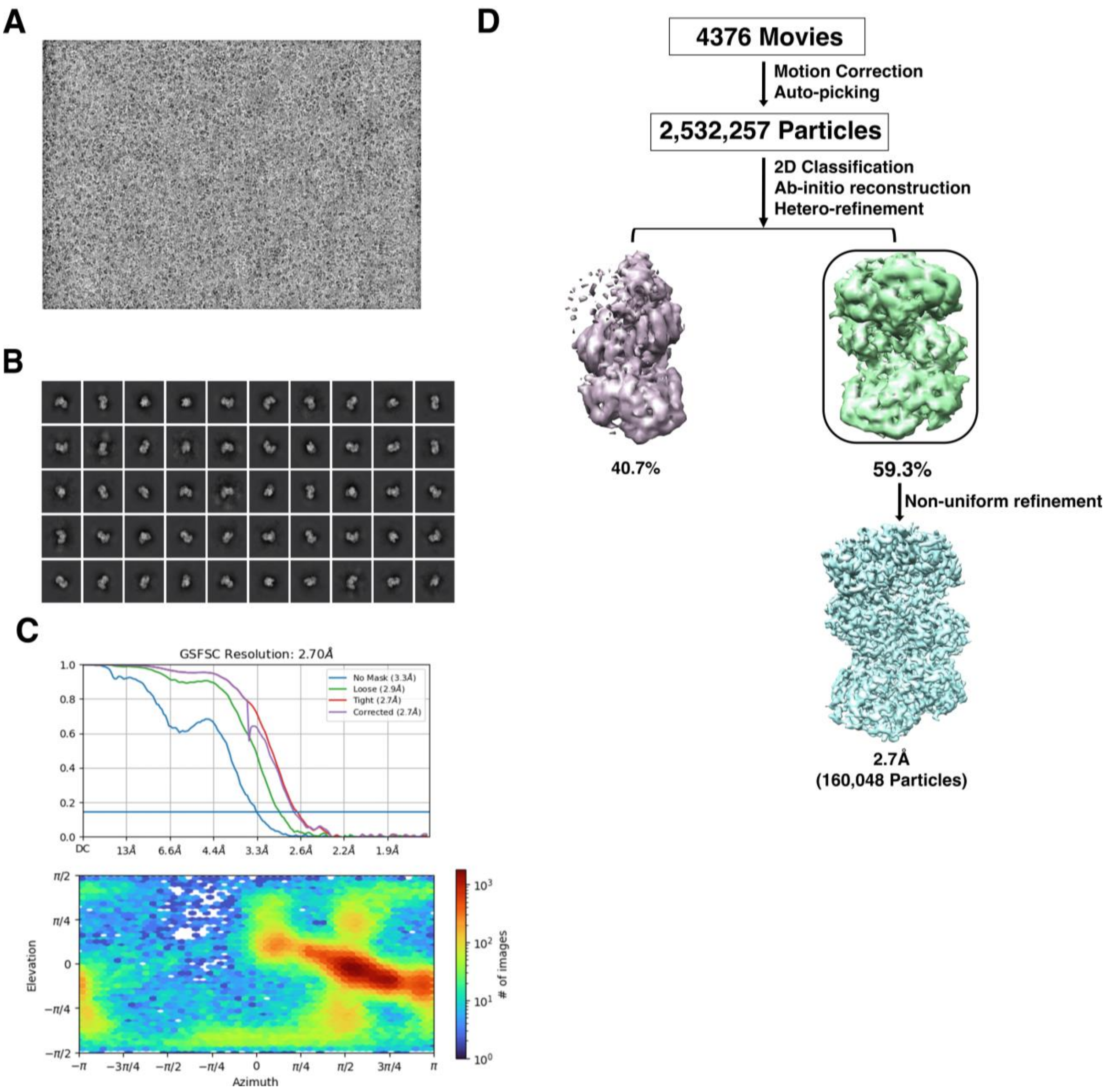


**Figure S1. Data processing flowchart for the single-particle cryo-EM analysis of the NB-protein apo complex.** (**A**) A representative motion-corrected micrograph of the NB-protein complex particles from 4376 movies. (**B**) Representative 2D CTF fit for the dataset. (**C**) Gold standard Fourier shell correlation (FSC) curve indicating overall nominal resolution at 2.7 Å using the FSC=0.143 criterion. (**D**) Workflow for Cryo-EM image processing of the NB-protein apo dataset.

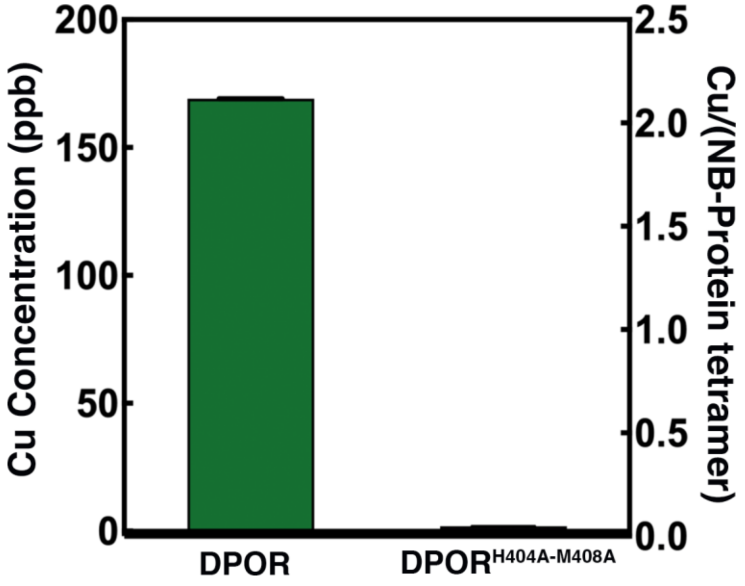


**Figure S2. Metal content in the wild-type and mutant DPOR complex samples.** The metal content was established based on an average of two measurements with the standard deviation less than 0.3%. In the wild-type NB grown in presence of Cu significant metal incorporation was observed. A stoichiometry of 2 Cu bound per NB-protein tetramer complex is observed. The Cu content was near the level of detection (LOD) for the mutant.

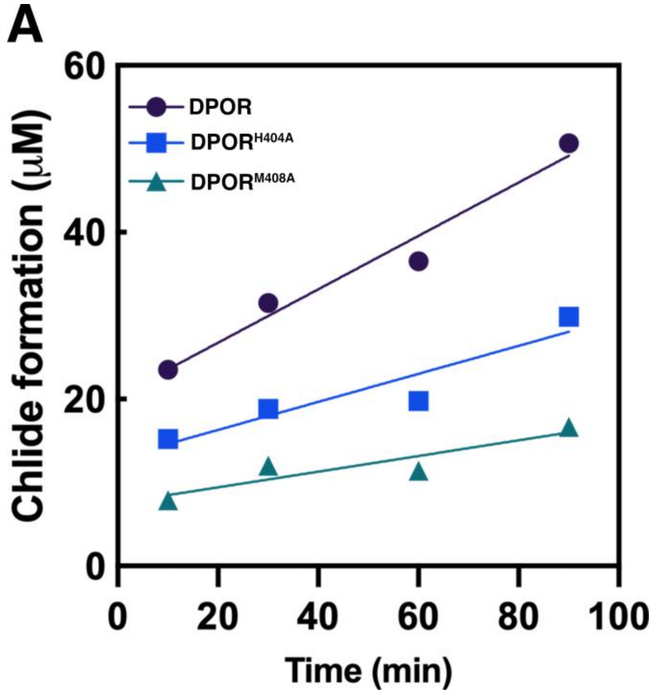


**Figure S3. Substitution of the Cu-coordinating residues results in perturbation of substrate reduction activity.** Pchlide reduction activity was measured for wild-type NB-protein and the BchN-BchB$^{H404A}$ or BchN-BchB$^{M408A}$ substituted complexes pre-bound to Pchlide. All experiments were carried out with BchL and initiated by adding ATP to the reaction. Time course of Chlide formation is shown. Alanine substitution of His-404 or Met-408 that coordinate the copper cluster reduce Pchlide reduction activity. Apparent rates for reduction of the bound Pchlide are: 0.32±0.02, 0.17±0.02, and 0.09±0.01 μM.min$^{-1}$ Chlide formed for wild-type, His-404A, and Met-408-A NB proteins, respectively. SE from n=3 experiments are denoted.

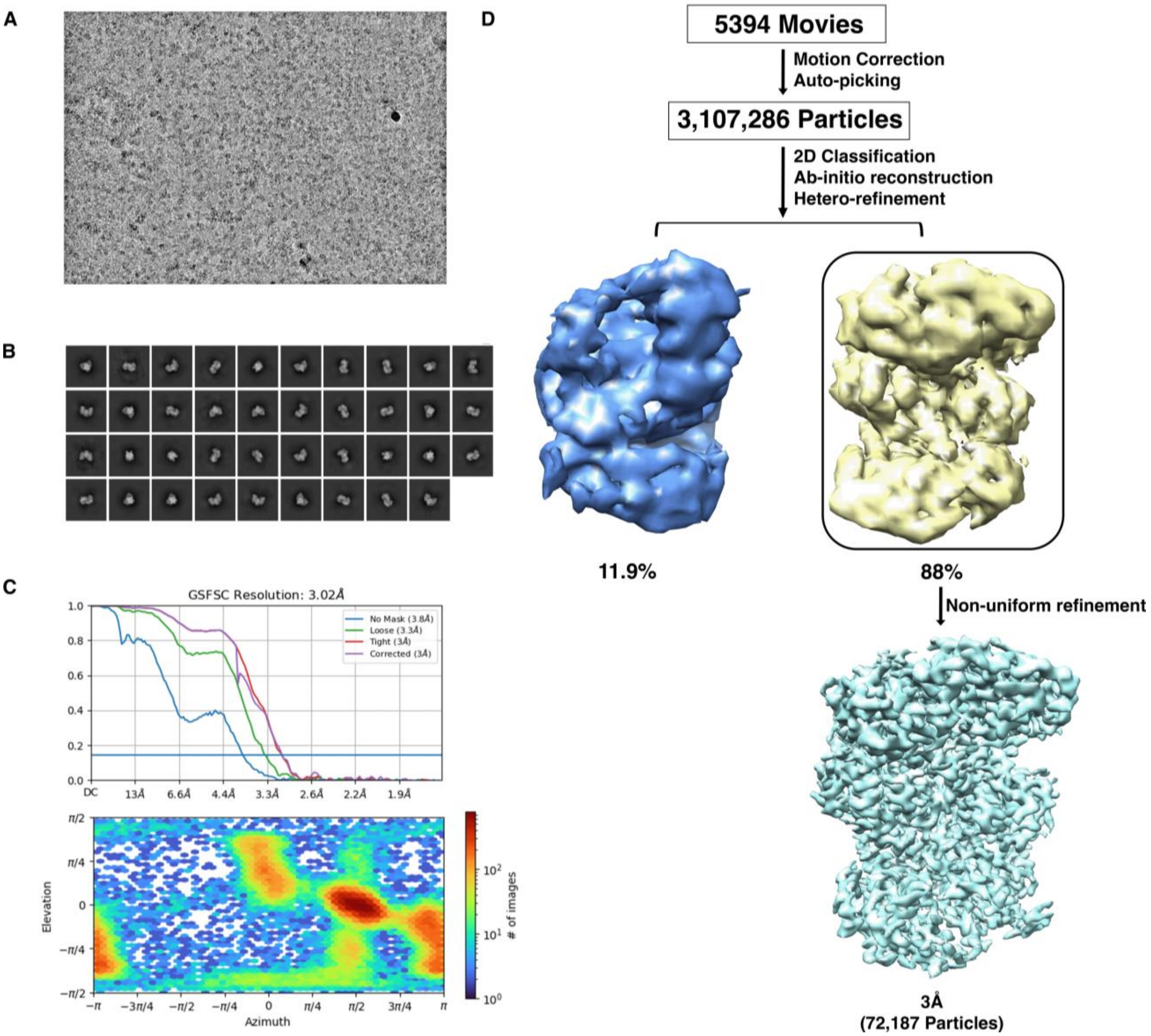


**Figure. S4. Data processing flowchart for the single-particle Cryo-EM analysis of the NB-protein-Pchlide complex.** (**A**) A representative motion-corrected micrograph of the NB-protein-Pchlide complex particles from 5394 movies. (**B**) Representative 2D CTF fit for the dataset. (**C**) Gold standard Fourier shell correlation (FSC) curve indicating overall nominal resolution at 3 Å using the FSC=0.143 criterion. (**D**) Workflow for Cryo-EM image processing of NB-protein-Pchlide complex.

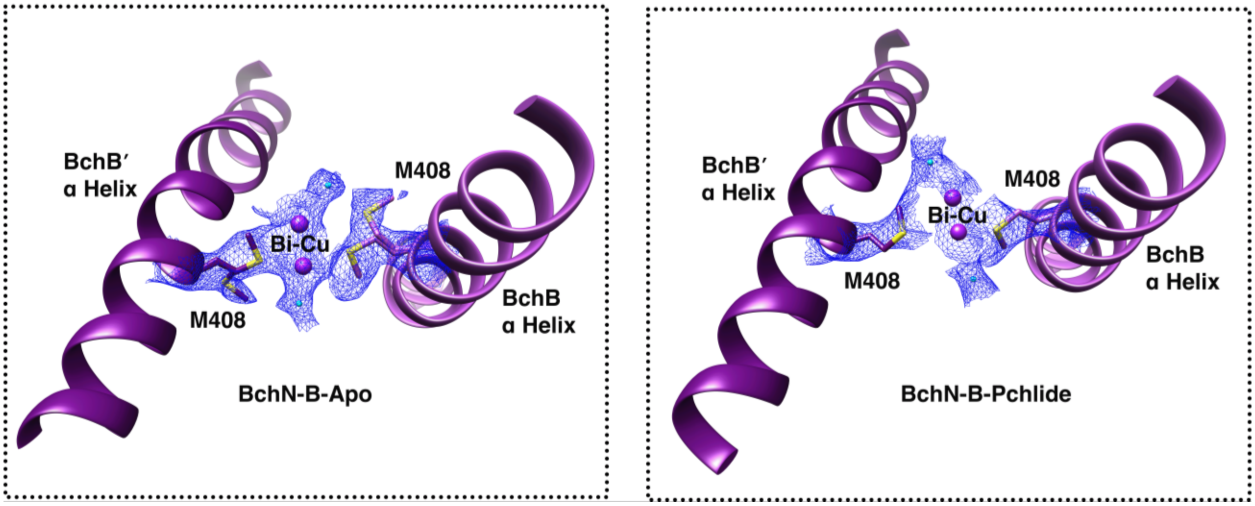


**Figure S5. Alternative side chain conformation of Met-404 in the NB-protein apo Cryo-EM structure** (**A**) Carved electron density depicting the dual side chain conformations for the Met-408 residue observed in the NB-protein apo Cryo-EM structure. (**B**) Carved electron density depicting the absence of the dual side chain conformation for Met-408 in the NB-protein-Pchlide bound complex. The (*) denotes the additional conformation observed for Met-408.

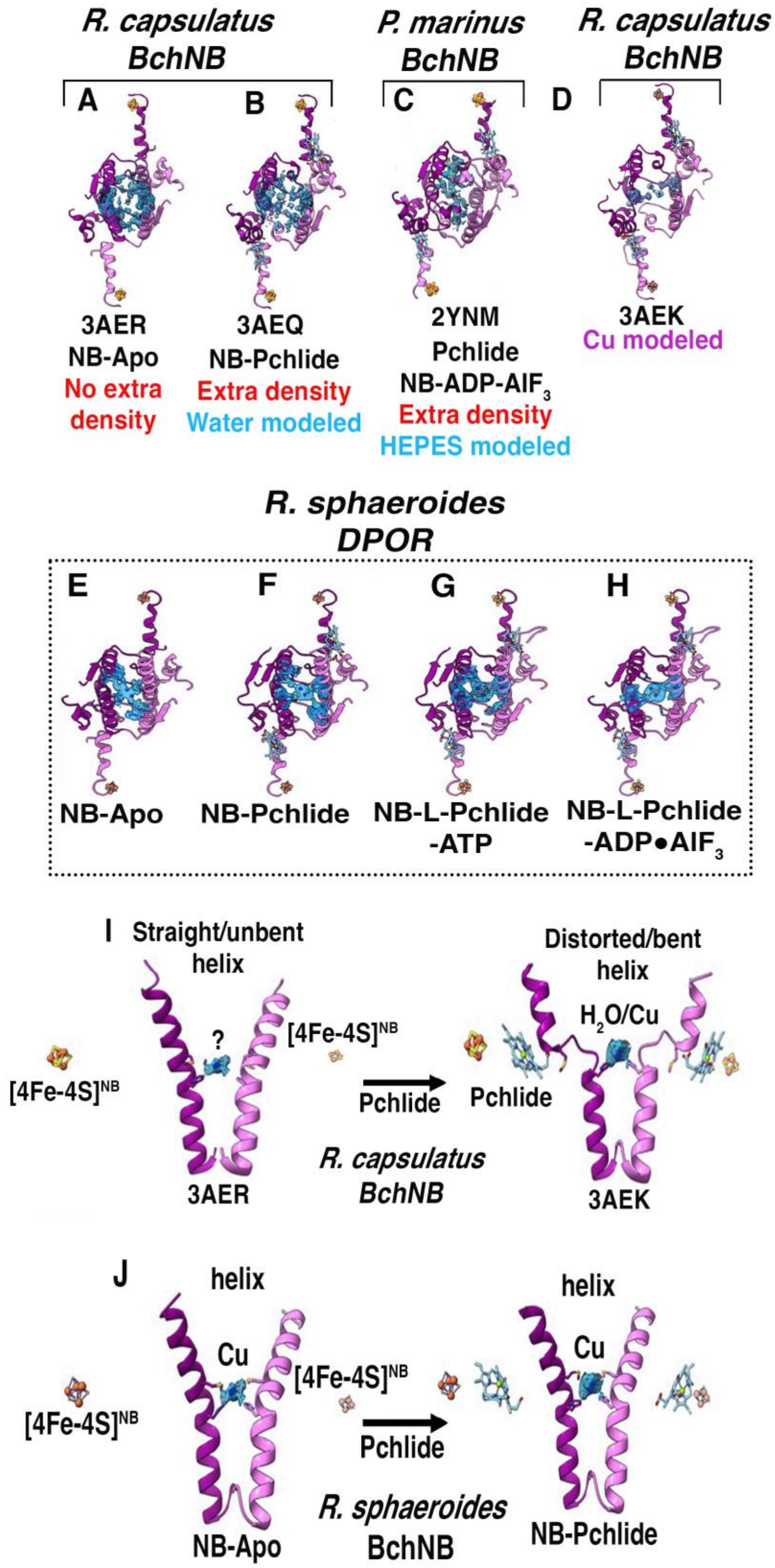


**Figure S6. Coordination of the Cu cluster or unresolved densities in the various structures of the NB-protein. (A-D)** Additional density found in various x-ray structures of the NB-protein are shown in blue and the two helices situated adjacently arise from each BchB subunit in the complex, and are colored pink and purple, respectively. The [4Fe-4S]$^{NB}$ and Pchlide molecules are also depicted. In the X-ray structures of (**A**) NB-protein apo and (**B**) Pchlide-bound NB-protein from *Rhodobacter capsulatus*, this density is either not modeled, or fit to water molecules. (**C**) In the *Prochlorococcus marinus* DPOR crystal structure, this density is modeled as two HEPES molecules. (**D**) Reanalysis of the original electron density map of the DPOR crystal structure (PDB: 3AEK) with copper modeled into the additional density shows an excellent fit for two Cu molecules coordinated by the His residues. (**E-H**) In all four of our Cryo-EM structures of *Rhodobacter sphaeroides* NB-protein we observe density for the bi-copper cluster. (**I**) In the crystal structures of *R. capsulatus* NB-protein, distortion of the two helices was observed upon Pchlide binding and thus proposed as a feature used to recognize substrate binding. (**J**) In our Cryo-EM structures of the NB-protein, we see a lack of structural changes in these helices in the absence or presence of Pchlide (or during turnover in the DPOR complex).

**A**

```
Conservation:                    9 9999999999 9999 99 9     999 9999999999999999  9999   99 9   9 999
BchB_from_R._sphaeroides     1   MKLTLWTYEGPPHVGAMRVATGMTGMHYVLHAPQGDTYADLLFTMIERRGKRPPVSYTTFQARDLGSDTA   70
BchB_from_R.capsulatus       1   MKLTLWTYEGPPHVGAMRVATAMKDLQLVLHGPQGDTYADLLFTMIERRNARPPVSFSTFEASHMGTDTA   70
BchB_from_P.marinus          1   MELTLWTYEGPPHIGAMRIATSMKGLHYVLHAPQGDTYADLLFTMIERRGSRPPVTYTTFQARDLGGDTA   70
Consensus_aa:                    McLTLWTYEGPPHlGAMRlATtMpshphVLHtPQGDTYADLLFTMIERRs.RPPVo@oTFpApchGsDTA
Consensus_ss:                           hh  hhhhhhhhh      eeeeee                       eeee   hhhhhh hh

Conservation:                     9       9  9  9 9  9    99999 99          9  9 9 9999 9   999 99 9 9
BchB_from_R._sphaeroides    71   ELFQSACRDAYERFQPQAIMVGSSCTAELIQDDTGGLADALSLPVPVVHLELPSYQRKENFGADESFLQI  140
BchB_from_R.capsulatus      71   ILLKDALAAAHARYKPQAMAVALTCTAELLQDDPNGISRALNLPVPVVPLELPSYSRKENYGADETFRAL  140
BchB_from_P.marinus         71   ELVKGHIFEAVERFKPEALLVGESCTAELIQDQPGSLAKGMGLNIPIVSLELPAYSKKENWGASETFYQL  140
Consensus_aa:                    bLhpshh..Ah.R@pPpAhhVt.oCTAELlQDpsstltcthsLslPlV.LELPtYp+KEN@GAsEoF..l
Consensus_ss:                    hhhhhhhhhhhhh    eeeee    hhhh   hhhhhhh       eeeee       hhhhhhhhhhh

Conservation:                     9 9                  9       9999   9999 999  9    9   99  9  99 9999
BchB_from_R._sphaeroides   141   CRKLARPM-------------ERTEKVSCNLLGPTALGFRHRDDILEVTRLLEGMGIAVNAVAPMGASPA  197
BchB_from_R.capsulatus     141   VRALAVPM-------------ERTPEVTCNLLGATALGFRHRDDVAEVTKLLATMGIKVNVCAPLGASPD  197
BchB_from_P.marinus        141   IRGLLKEISEDSSNNAKQSWQEEGRRPRVNLLGPSLLGFRCRDDVLEIQKILGENGIDINVIAPLGASPS  210
Consensus_aa:                    hR.Lh..h.............Ecs.csphNLLGsohLGFRhRDDlhElp+lL...GI.lNhhAPhGASPs
Consensus_ss:                    hhhh                          eeee         hhhhhhhhhhhh     eeeee      hh

Conservation:                    9   9  9  99  999  99      9   9 9 99 99999 99 99  9     9
BchB_from_R._sphaeroides   198   DIARLGAAHFNVLLYPETGESAARWAEKTLKQPYTKTVPIGVGATRDFVAEVAALAGVAP---VADDSRL  264
BchB_from_R.capsulatus     198   DLRKLGQAHFNVLMYPETGESAARHLERACKQPFTKIVPIGVGATRDFLAEVSKITGLPV---VTDESTL  264
BchB_from_P.marinus        211   DLMRLPKADANVCLYPEIAESTCLWLERNFKTPFTKVVPIGVKATQDFLEELYELLGMEVSNSISNSDQS  280
Consensus_aa:                    Dl.+Ls.AchNVhhYPEhtESht.@..lhGh.s...lsspsp.
Consensus_ss:                    hhhh     eeeee hhhhhhhhhhhhhh    eeee      hhhhhhhhhhhhhhh    h   hhhhhhh

Conservation:                      99 9 9999 99999999 999 999 9999 9  9 999999 9 9 99 99  9 99   99999
BchB_from_R._sphaeroides   265   RQPWWSASVDSTYLTGKRVFLFGDATHVIAAARVARDEMGFEVVGMGCYNREFARPMRAAAKGYGLEALV  334
BchB_from_R.capsulatus     265   RQPWWSASVDSTYLTGKRVFIFGDGTHVIAAARIAAKEVGFEVVGMGCYNREMARPLRTAAAEYGLEALI  334
BchB_from_P.marinus        281   KLPWYSKSVDSNYLTGKRVFIFGDGTHVLAAARIANEELGFEVVGIGTYSREMARKVRAAATELGLEALI  350
Consensus_aa:                    +bPW@S.SVDSsYLTGKRVFlFGDtTHVlAAARlA.cEhGFEVVGhGhYsREhAR.hRhAA..hGLEALl
Consensus_ss:                    hhhhhhhh  hhhh    eeeee      hhhhhhhhhhhh   eeeeeee     hhhhhhhhhhh    eeee

                                                                                                 *His
Conservation:                    9 999999   9   9999 9999999  99 99 999999 9 9999 9999 9999 99999 99 9 9
BchB_from_R._sphaeroides   335   TDDYLEVEEAIQALAPELILGTQMERHIAKRLGIPCAVISAPVHVQDFPARYSPQMGFEGANVLFDTWIH  404
BchB_from_R.capsulatus     335   TDDYLEVEKAIEAAAPELILGTQMERNIAKKLGLPCAVISAPVHVQDFPARYAPQMGFEGANVLFDTWVH  404
BchB_from_P.marinus        351   TNDYLEVEESIKECAPELVLGTQMERHSAKRLGIPCAVISTPMHVQDVPARYSPQMGWEGANVIFDDWVH  420
Consensus_aa:                    TsDYLEVEctIp.hAPELlLGTQMERp.AK+LGlPCAVIShPhHVQDhPARYtPQMG@EGANVlFDsWlH
Consensus_ss:                    e  hhhhhhhhhh    eeee    hhhhhhh     eeeee  eee          ee hhhhhhhhhhhhh

                                    * Met
Conservation:                    99 9999999  999 9999 9       9 99 9                                  9
BchB_from_R._sphaeroides   405   PLTMGLEEHLLTMFREDFEFHD--EAGPSHHGGKAVPASAPRADEAAEALPATGAETAEGGSIPPEAVPP  472
BchB_from_R.capsulatus     405   PLVMGLEEHLLTMFREDFEFHD--AAGASHHGGKAV------AREESPVAPADLAPAATSDT--PAAPSP  464
BchB_from_P.marinus        421   PLMMGLEEHLIGMFRHDFEFTDGHQSHLGHLGGHA------------------SETKTSSK--GINQSP  469
Consensus_aa:                    PLhMGLEEHLlsMFRcDFEFhD...t..tHhGG+A...................t.h.ptsp..s.s.sP
Consensus_ss:                    hhhhhhhhhhhhhh

Conservation:                        9      9 9     9 99 999999999  9999  9    9    9  999   9999
BchB_from_R._sphaeroides   473   AAAAAAEAPAGEIVWLTDAERELKKIPFFVRGKARRNTEKFAAEKGLTRISIETLYEAKAHYAR  536
BchB_from_R.capsulatus     465   VVVTQA---SGEIRWMPEAERELRKIPFFVRGKAKRNTELYAAHKGVCDITVETLYEAKAHYAR  525
BchB_from_P.marinus        470   NNHSPA---GESIHWTSEGESELAKIPFFVRGKVRRNTEKYARQAGCREIDGETLLDAKAHFGA  530
Consensus_aa:                    sshs.A...t.pI.Whs-tEpEL.KIPFFVRGKh+RNTEb@A.p.Gh.cIs.ETLh-AKAH@t.
Consensus_ss:                                    hhhhhhhh    hhhhhhhhhhhhhhhhh        hhhhhhhhhhh
```

**Figure S7. Sequence alignment of BchB highlighting the conservation of Met-404 and His-408.** (**A**) An amino acid sequence alignment of BchB from *R. sphaeroides*, *R. capsulatus* and *P. marinus* is shown. His-404 and Met-408 are conserved across the three organisms.

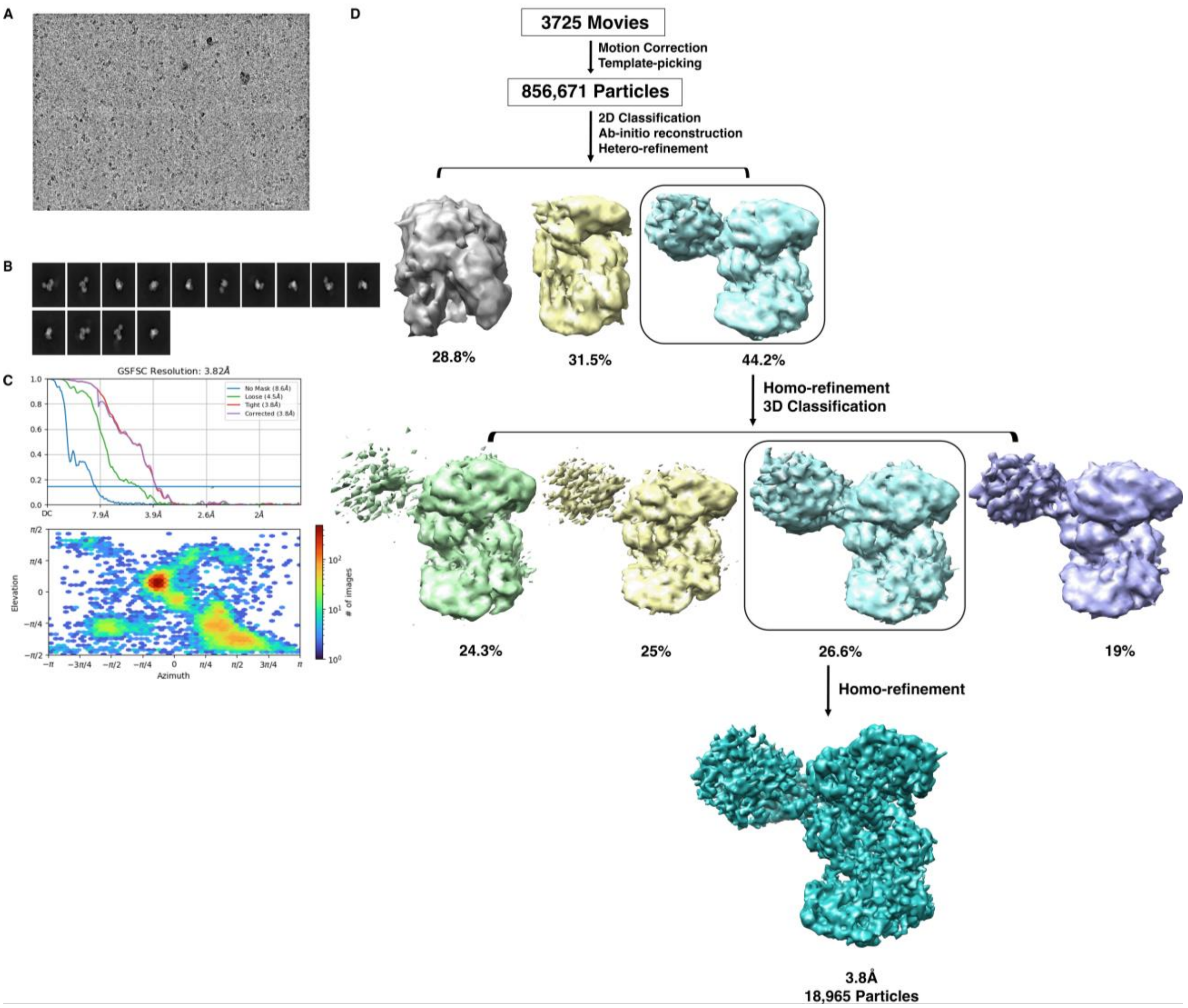


**Figure S8. Data processing flowchart for the single-particle Cryo-EM analysis of the DPOR complex under turnover.** (**A**) A representative motion-corrected micrograph of the DPOR complex under turnover (in the presence of ATP) particles from 3725 movies. (**B**) Representative 2D CTF fit for the dataset showing 1:1 complex of NB-protein:L-protein. (**C**) Representative 2D CTF fit for the dataset showing the L-proteim alone. (**D**) Representative 2D CTF fit for the dataset showing the NB-protein alone. (**E**) Gold standard Fourier shell correlation (FSC) curve indicating overall nominal resolution at 3.8 Å using the FSC=0.143 criterion. (**F**) Workflow for Cryo-EM image processing of DPOR under turnover complex.

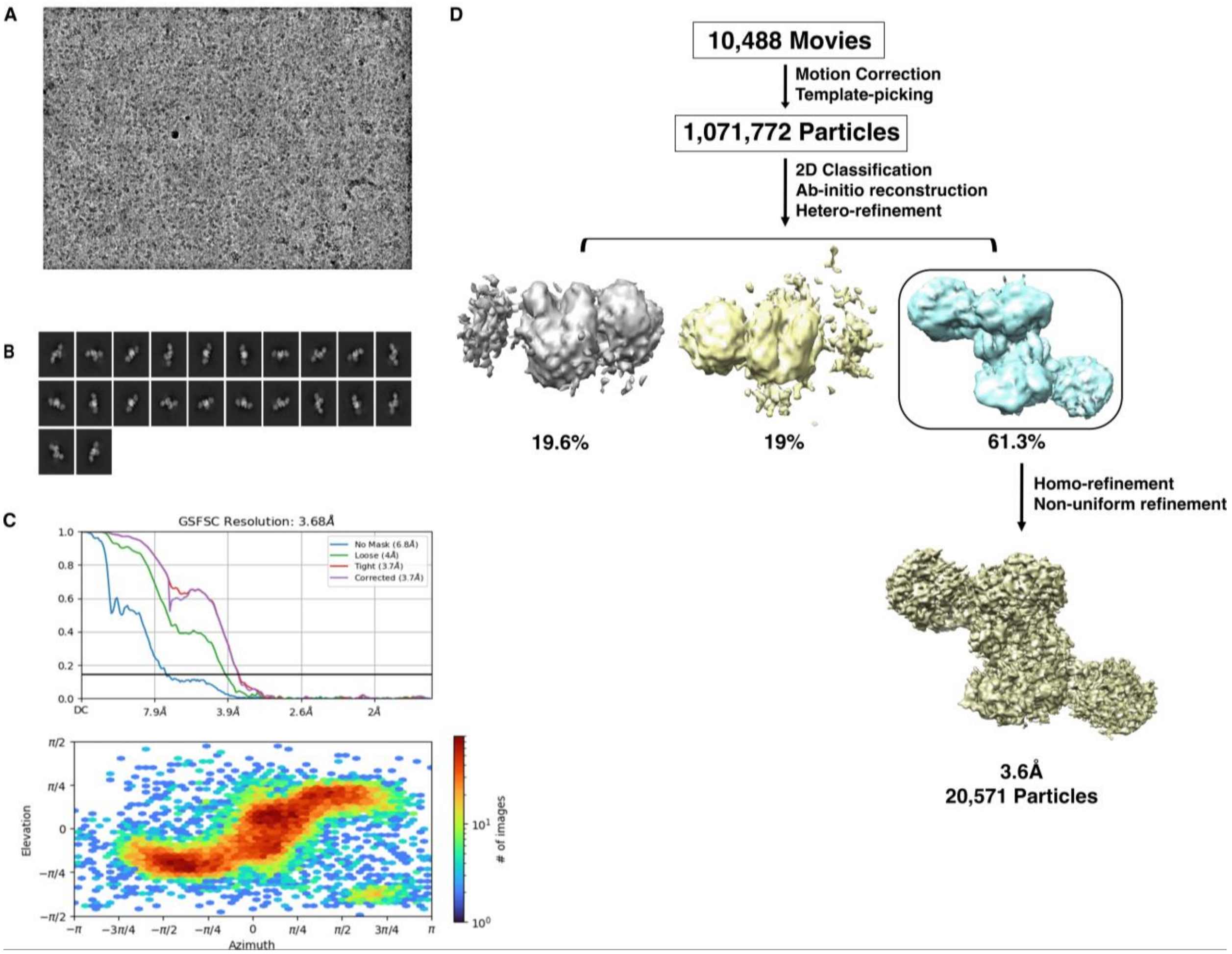


**Figure S9. Data processing flowchart for the single-particle cryo-EM analysis of DPOR in the presence of ADP-AlF$_3$ complex.** (**A**) A representative motion-corrected micrograph of the DPOR transition-state complex in the presence of ADP-AlF$_3$ complex particles from 10,488 movies. (**B**) Representative 2D CTF fit for the dataset. (**C**) Gold standard Fourier shell correlation (FSC) curve indicating overall nominal resolution at 3.6 Å using the FSC=0.143 criterion. (**D**) Workflow for cryo-EM image processing of DPOR complex in the presence of ADP-AlF$_3$.

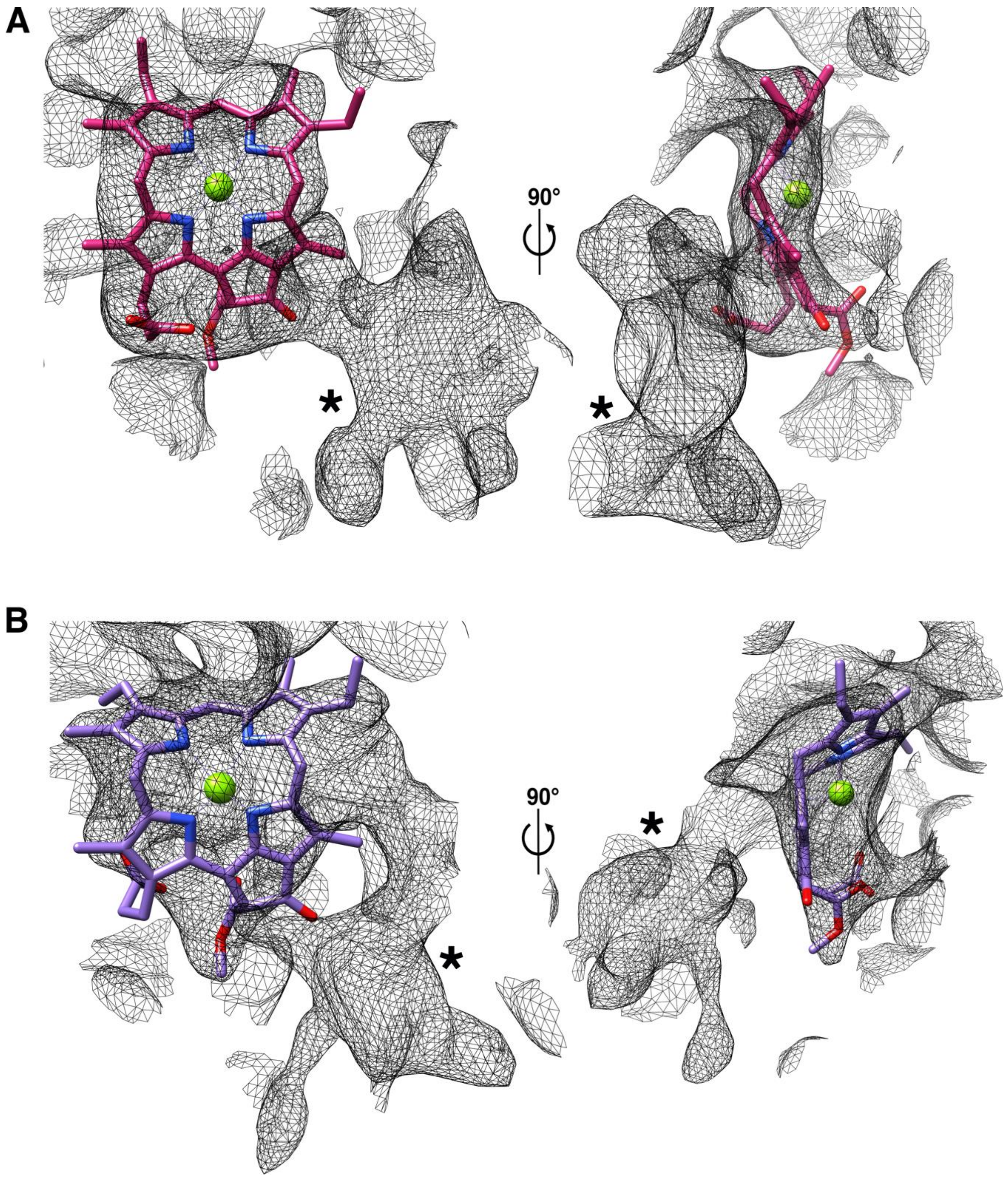


**Figure S10. Extra density near the Pchlide in the DPOR transition-state CryoEM structures. A)** BchN-BchB or **B)** BchN′-BchB′ active sites in the DPOR-transition-state CryoEM structure shows extra density marked with * near the Pchlide. This density is sandwiched between the Pchlide and the C-terminal of BchB (in-trans). This figure was generated at contour level 0.114 with the zone radius as 4.

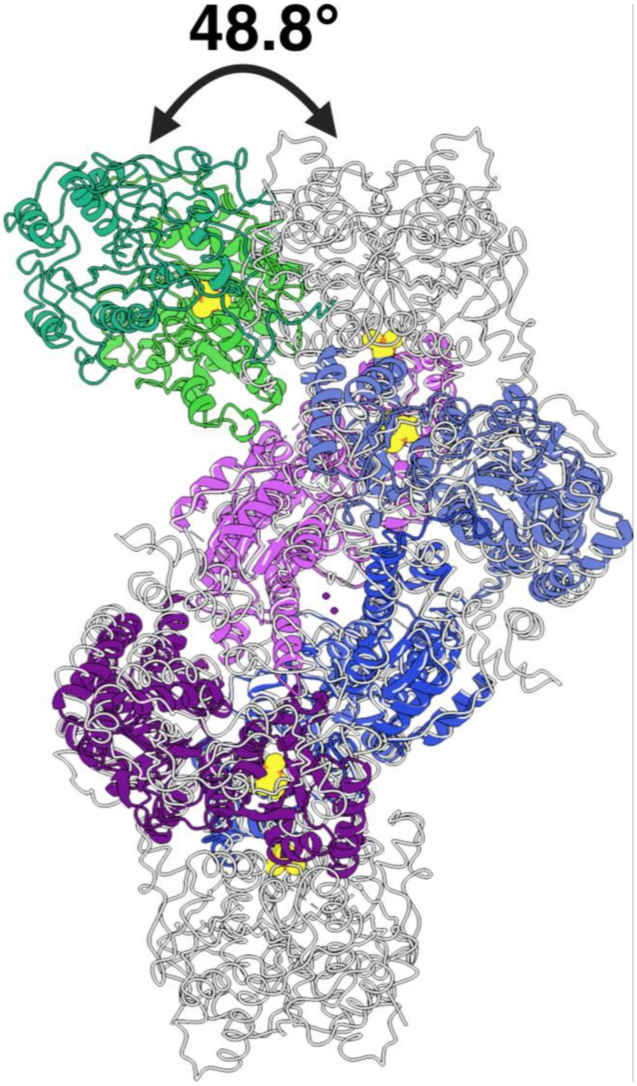


**Figure S11. Rolling-motion of the L-protein on the NB-protein in the CryoEM turnover complex of DPOR.** Superposition of the *P. marinus* DPOR crystal structure solved in the presence of ADP-$AlF_3$ (PDB: 2YNM in grey) with the CryoEM structure of *R. sphaeroides* DPOR (colored subunits; under turnover conditions) shows a large 49º difference in the docking positions of the respective L-proteins on the NB-protein.

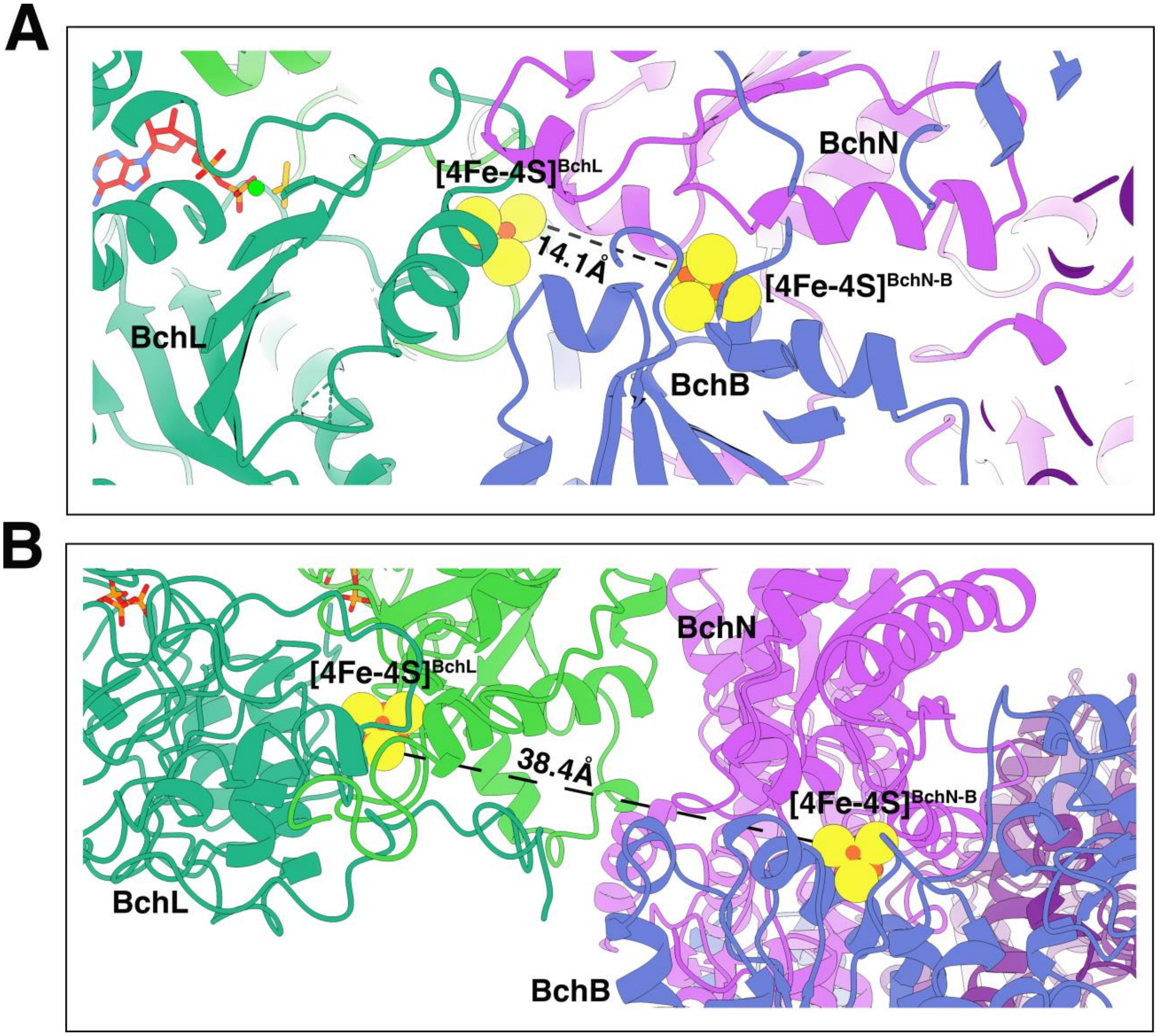


**Figure S12. Rolling motion of the L-protein on the NB-protein results in changes in intramolecular distance between the FeS clusters.** Comparison of the distances between the $[4Fe\text{-}4S]^{L}$ and $[4Fe\text{-}4S]^{NB}$ clusters in the ADP-$AlF_3$ bound crystal structure (PDB: 2YNM) and the cryo-EM structure of the DPOR complex under turnover. (**A**) The distance between the $[4Fe\text{-}4S]^{L}$ and $[4Fe\text{-}4S]^{NB}$ clusters in the ADP-$AlF_3$ bound crystal structure is 14.1Å. (**B**) The distance between the two clusters in the Cryo-EM structure of DPOR under turnover condition (in the presence of ATP) is 38.4Å.

**Supplementary Table 1.** Cryo-EM data collection and model statistics of NB-protein complex and NB-complex bound to Pchlide.

| | NB-Protein (Apo)<br>(EMDB-43443)<br>(PDB-8VQH) | NB-Protein (Pchlide bound)<br>(EMDB-43444)<br>(PDB-8VQI) |
|---|---|---|
| **Data Collection** | Titan Krios GATAN K3 BioQuantum | Titan Krios GATAN K3 BioQuantum |
| Magnification | 105,000 | 105,000 |
| Voltage (kV) | 300 | 300 |
| Spherical Aberration (mm) | 2.7 | 2.7 |
| Electron Exposure ($e^-$/ Å$^2$) | 50 | 50 |
| Defocus range (μm) | -0.1 to -2.4 | -0.1 to -2.4 |
| Pixel size (Å, Physical/Digital) | 0.825 | 0.825 |
| Energy Filter Slit Width (eV) | 15 | 15 |
| Movies | 4376 | 5394 |
| **Map Statistics and Post-Processing** | | |
| Symmetry imposed | C1 | C1 |
| Map Resolution (Å) | 2.7 | 3 |
| FSC threshold | 0.143 | 0.143 |
| **Model Composition** | | |
| Chains | 8 | 4 |
| Non-hydrogen atoms | 12659 | 12573 |
| Protein residues | 1645 | 1643 |
| Ligands | SF4:2<br>CU:2 | SF4:2<br>CU:2<br>PMR:2 |
| Water | 2 | 2 |

| | | |
|---|---|---|
| Bonds (RMSD)<br>Length (Å) (#>4σ)<br>Angles (º) (#>4σ) | <br>0.013 (13)<br>2.008 (107) | <br>0.012 (0)<br>1.699 (33) |
| **Validation** | | |
| Molprobity Score | 3.13 | 2.98 |
| All-atom clash score | 18.09 | 11.35 |
| Ramachandran plot<br>Favored (%)<br>Allowed (%)<br>Outliers (%) | <br>96.64<br>3.18<br>0.18 | <br>96.57<br>3.43<br>0.00 |

**Supplementary Table 2.** Cryo-EM data collection and model statistics for the DPOR complexes solved under turnover (ATP) or transition-state (ADP-$AlF_3$) conditions.

| | DPOR under turnover (ATP)<br>(EMDB-43446)<br>(PDB-8VQJ) | DPOR under transition state (ADP-$AlF_3$) |
|---|---|---|
| **Data Collection** | Titan Krios GATAN K3 BioQuantum | Titan Krios GATAN K3 BioQuantum |
| Magnification | 105,000 | 105,000 |
| Voltage (kV) | 300 | 300 |
| Spherical Aberration (mm) | 2.7 | 2.7 |
| Electron Exposure ($e^-$/ $Å^2$) | 50 | 50 |
| Defocus range (μm) | -0.1 to -2.4 | -0.1 to -2.4 |
| Pixel size (Å, Physical/Digital) | 0.825 | 0.825 |
| Energy Filter Slit Width (eV) | 15 | 15 |
| Movies | 3725 | 10,488 |
| **Map Statistics and Post-Processing** | | |
| Symmetry imposed | C1 | C1 |
| Map Resolution (Å) | 3.7 | 3.6 |
| FSC threshold | 0.143 | 0.143 |
| **Model Composition** | | |
| Chains | 10 | 12 |
| Non-hydrogen atoms | 17018 | 20663 |
| Protein residues | 2233 | 2710 |
| Ligands | SF4:2<br>CU:2<br>MG:2<br>ATP:1<br>ADP:1 | SF4:4<br>CU:2<br>MG:2<br>PMR:2<br>ADP.AlF3: 4 |

| | | |
|---|---|---|
| Water | 0 | 0 |
| Bonds (RMSD)<br>Length (Å) (#>4σ)<br>Angles (º) (#>4σ) | <br>0.010 (21)<br>1.533 (149) | <br>0.013(24)<br>1.955(148) |
| **Validation** | | |
| Molprobity Score | 3.13 | 4.10 |
| All-atom clash score | 29.78 | 39.74 |
| Ramachandran plot<br>Favored (%)<br>Allowed (%)<br>Outliers (%) | <br>90.86<br>9.00<br>0.14 | <br>77<br>22.4<br>0.67 |
